# Identification of Multiple Prognostic Biomarker Sets for Risk Stratification in SKCM

**DOI:** 10.1101/2025.02.27.640548

**Authors:** Shivani Malik, Ritu Tomer, Akanksha Arora, Gajendra P. S. Raghava

## Abstract

Most of the existing studies have identified a single profile of prognostic biomarkers for predicting high-risk cancer patients using transcriptomics data. In this study, we propose multiple distinct sets of prognostic biomarkers for predicting high-risk Skin Cutaneous Melanoma (SKCM) patients. Our primary analysis reveals that the expression of certain genes, such as CREG1, PCGF5, and VPS13C, strongly correlates with overall survival (OS) in SKCM patients. We developed machine learning-based prognostic models to predict 1-, 3-, and 5-year overall survival using gene expression profiles. State-of-the-art feature selection techniques were employed to identify the first prognostic biomarkers set consisting of 20 genes. Machine learning models built to predict high-risk patients using this set of biomarkers and achieved an AUC of 0.90 and a Kappa of 0.58 in distinguishing high-risk SKCM patients. Similarly, a second independent set of 20 prognostic genes was identified, with no overlap with the first set. The best model trained on this second set achieved an AUC of 0.89 with a Kappa of 0.56, while the fifth biomarker set achieved the highest performance with an AUC of 0.91 and a Kappa of 0.64. This process was repeated to obtain a total of seven distinct prognostic biomarker sets, each containing 20 unique genes. The predictive performance of these models varied between 0.84 and 0.91 for AUC and 0.48 to 0.64 for Kappa on the validation dataset. These findings demonstrate that it is possible to identify multiple independent sets of prognostic biomarkers, each capable of accurately predicting high-risk SKCM patients.

**Highlights:**

1. Prediction of high-risk SKCM patients from transcriptomics data
2. Identification of multiple set of prognostic biomarkers
3. Identification of set of biomarkers using feature selection technique
4. Development of machine learning based models for prediction
5. Pathway analysis of selected set of biomarker genes

## Introduction

Skin Cutaneous Melanoma accounts for around 3–5% of all malignancies and is a very aggressive tumor that arises from melanocyte cells and primarily affects the skin. However, it can also occur in mucous membranes and internal organs (Sinagra & Sciume, 2020; F. Wang et al., 2022). The third most frequent type of skin cancer, skin cutaneous melanoma, affects 6.8% to 20% of cases (Z. Wang & Liu, 2021). According to projections, there will be 2,001,140 new cases of cancer and 611,720 cancer-related fatalities in the US in 2024. Additionally, there will likely be over 8,000 deaths and 100,640 new cases of SKCM (Siegel et al., 2024). Globally, the incidence of melanoma continues to rise, driven primarily by environmental factors, particularly excessive ultraviolet (UV) radiation exposure, which plays a central role in melanoma carcinogenesis (Tracey & Vij, 2019). Human skin color is primarily determined by the amount of melanin, with darker skin containing larger melanocytes that produce more melanin, offering protection against UV radiation (Aoude et al., 2014; Kaidbey et al., 1979).

SKCM can originate from benign melanocytic nevi, including dysplastic, acquired, or congenital forms, or may develop independently. In recent years, both its prevalence and mortality rates have risen significantly. Surgical excision remains the most effective treatment, highlighting the importance of early diagnosis and intervention (Christodoulou et al., 2021; Kardynal & Olszewska, 2014; Merighi et al., 2009; Seitz et al., 2021). Advances in genomic analysis and high-throughput technologies have provided deeper insights into melanoma biology, improving detection and treatment strategies (Scatena et al., 2021). The malignant transformation of melanocytes is driven by genetic alterations, particularly in the BRAF and NRAS genes (Guan et al., 2015). These mutations are central to melanoma development and progression, influencing tumor behavior and patient response to therapies (Garman et al., 2017). Despite significant progress in uncovering the genetic underpinnings of melanoma, late-stage diagnosis remains a major obstacle, limiting the effectiveness of treatments such as surgery, immunotherapy, and targeted therapies (Y. He & Wang, 2023).

Recent studies have focused on identifying invasion-related biomarkers for individualized treatment and prognosis prediction in SKCM. One such study analyzed 124 invasion-associated genes (IAGS) and selected 20 prognostic genes to develop a gene expression-based model (E. Yang et al., 2023). Similarly, another study used transcriptome data from TCGA to find a predictive signature based on m6A-related lncRNAs (Lin et al., 2024). Another study developed a 10 ferroptosis-related gene (FRG) signature for predicting overall survival in SKCM (Ping et al., 2022). Additionally, eight immune-related lncRNA signatures were identified as a prognostic model for melanoma and validated using Kaplan-Meier analysis (Xiao et al., 2021). In recent studies, a prognostic signature comprising seven NLR-related genes was identified using LASSO-Cox analysis. This signature effectively stratified SKCM patients into high and low-risk groups, demonstrating a strong correlation with overall survival outcomes. Additionally, it revealed distinct immune microenvironment patterns characterized by the activation of inflammatory and interferon pathways and showed potential for predicting responses to immune checkpoint blockade therapy (Geng et al., 2023). A chemokine-related 14-gene prognostic model was developed, revealing distinct immune characteristics and stratification between low- and high-risk groups. This model demonstrates potential in predicting immunotherapy outcomes and chemotherapy efficacy and aiding clinical decision-making for SKCM patients (X. Ding et al., 2023). While these studies provide valuable insights, they typically focus on a limited set of biomarkers within a specific context, as shown in the **Supplementary Table S1**.

Previous studies have mostly focused on pathway-related biomarkers. To the best of our knowledge, these methods typically identify only a single set of biomarkers. Moreover, they usually select prognostic biomarkers from a limited subset of data, such as genes related to invasion, immune response, or lncRNAs. In this study, we take a novel approach by attempting to generate biomarker sets from all available genes rather than focusing on just a subset. Additionally, we select a secondary set from the remaining genes after identifying an initial set of biomarkers. This process is repeated, leading to the identification of seven distinct sets of prognostic biomarkers.

## Methodology

### Data Collection

Transcriptomic profiling and clinical data were retrieved from The Cancer Genome Atlas (TCGA), a major cancer research database (C. Yan et al., 2022) using the TCGAbiolinks package from BiocManager in R. The retrieved expression data, which included 473 samples and 60,660 genes, was normalized to Transcripts Per Kilobase Million (TPM) values before downstream analysis using R script.

### Preprocessing of Data

We first removed duplicate features and filtered out genes that contained more than 50% zero values. Next, we used the caret package in R to remove low-variance features, ensuring that only informative genes were retained. After preprocessing, we retained 23,009 genes for further analysis. Upon merging this gene expression data with the corresponding clinical data, we identified a cohort of 287 patients with available survival information.

#### Z-score scaling

In order to ensure that each feature has a mean of 0 and a standard deviation of 1, this transformation standardizes the data by subtracting the mean of each feature and dividing it by its standard deviation. By standardizing the data size, this technique makes it possible to compare features more accurately (Tihagam & Bhatnagar, 2023).

## Statistical Methods

### Correlation analysis

The associations between gene expression and OS time were evaluated using Pearson correlation analysis in Base R. Genes with a p-value < 0.05 were considered significant, providing insights into their potential impact on SKCM patient survival (C. Ma & Xie, 2024). Positively correlated genes were further analyzed using Gene Ontology (GO) (J. Wang & Yang, 2022) and Reactome pathway analysis (Fabregat et al., 2017) to identify enriched biological processes and pathways relevant to melanoma progression, utilizing the gseapy package in Python.

### Survival Analysis

Patients were divided into high- and low-risk groups based on the median overall survival (OS) time. Univariate Cox regression analysis was conducted using the Cox proportional hazards model (R.-H. Yang et al., 2021). Significant genes were selected based on p-values < 0.05 using the survival package in R. Subsequently, LASSO regression analysis (Huang et al., 2022) was performed using the glmnet package in R to identify prognostic genes, with an optimal lambda value of 0.03. Depending on the formula:

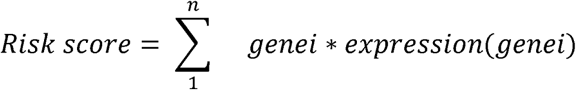

Identified prognostic genes were then analyzed using the Drug Gene Interaction Database (DGIdb) (Cannon et al., 2024) to predict potential therapeutic agents based on known drug-gene interactions.

### AI-based Methods

We employed various feature selection techniques to identify relevant genes for the prognostic model. Overall survival time was categorized into four classes: Class 0 (0–1 years), Class 1 (1–3 years), Class 2 (3–5 years), and Class 3 (>5 years).

### Feature Selection Techniques

To identify the most relevant features for the prognostic model, we applied several established feature selection methods, including Linear Support Vector Classification with L1 penalty (SVC-L1), Recursive Feature Elimination (RFE), Sequential Feature Selection (SFS), and SelectKBest. These methods have been successfully employed in similar studies (Jiang et al., 2023; Kamkar et al., 2016; S. Ma & Huang, 2008)

### Identification of the Primary Set of Biomarkers

We utilized a variety of feature selection techniques, starting with the Recursive Feature Elimination algorithm (Darst et al., 2018), which recursively ranks features based on their relevance and selects the optimal subset. In the second method, SelectKBest, we applied the f_classif evaluation function to identify the K most relevant features. We also used Linear Support Vector Classification with L1 regularization, which selects features by penalizing less important ones, achieving sparsity, and improving model performance (Bhalla et al., 2019). Specifically, we obtained the absolute values of SVC-L1 coefficients and used the mean coefficient value as a threshold to select features with importance scores above the mean. Sequential Feature Selection was employed to add or remove features to optimize performance sequentially. These techniques helped identify a robust initial biomarker set for further analysis using the Scikit-learn package (Pedregosa, F. and Varoquaux, G. and Gramfort, A. and Michel et al., 2011).

### Identification of Additional Biomarker Sets

For the secondary biomarker set, features identified in the primary set were excluded before applying the SVC-L1 method to ensure the independence of the sets. In the case of the third biomarker set, invasion-associated genes, along with features previously identified through RFE and SVC-L1 in earlier analyses, were systematically removed prior to the selection process. This process is visualized in **Figure 3**, where the Venn diagram highlights the common and distinct features across the biomarker sets and invasion-associated genes. This approach facilitated the identification of non-overlapping and distinct biomarker sets for further analysis.

### ML Models

Class imbalance is addressed by using the Synthetic Minority Over-sampling Technique (SMOTE). To evaluate the model’s robustness and performance, stratified five-fold cross-validation was conducted using an 80:20 train set to test set ratio. Multi-Layer Perceptron (MLP), Decision Tree (DT), Extreme Gradient Boosting (XGB), Support Vector Machine (SVM), Extra Trees Classifier (ET), Random Forest (RF), k-Nearest Neighbors (KNN), Light Gradient Boosting Machine (LightGBM), Categorical Boosting (CatBoost), Adaptive Boosting (AdaBoost), and Logistic Regression (LR) are some of the machine learning (ML) algorithms that we used. We have built prediction models using these ML algorithms on different sets of features.

### Ensemble Models

We used a voting classifier by utilizing multiple base classifiers, i.e., RF, ET, and LightGBM, to make individual predictions. We used hard voting, where the majority of the votes from the classifiers determined the final class prediction. This method helps aggregate the strengths of different models to achieve better generalization. We also applied a stacking classifier and trained several different base models, including ET & RF. The predictions of these base models were used as inputs to LR as a meta-model, which was trained to combine the base model outputs and make the final prediction. We tried catBoost as a meta-model & RF, ET, and XGB as base models. This two-level architecture leverages the strengths of multiple models by allowing the meta-model to learn how to best combine their predictions.

## Results

### Correlation Analysis

Our univariate Cox regression analysis identified 4,324 genes significantly associated with overall survival time. We selected the top genes with the most significant positive and negative correlations, as shown in **Table 1**. Specifically, the genes CREG1 and PCGF5 exhibited positive correlations with OS with correlation coefficients of 0.40 and 0.38, respectively. Conversely, the genes AC008687.4 and TTYH2 showed negative correlations with OS, with coefficients of −0.23 and −0.23, respectively. A positive correlation suggests that higher expression levels of these genes are associated with longer survival, whereas a negative correlation indicates that higher expression levels are linked to poorer survival outcomes. The Pearson correlation coefficient (r) was used to quantify these associations.

**Table 1.**
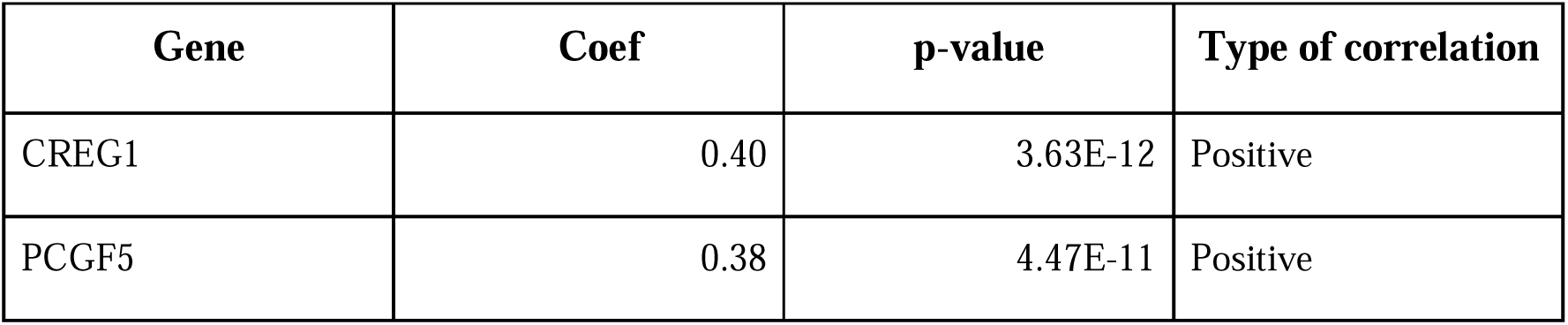
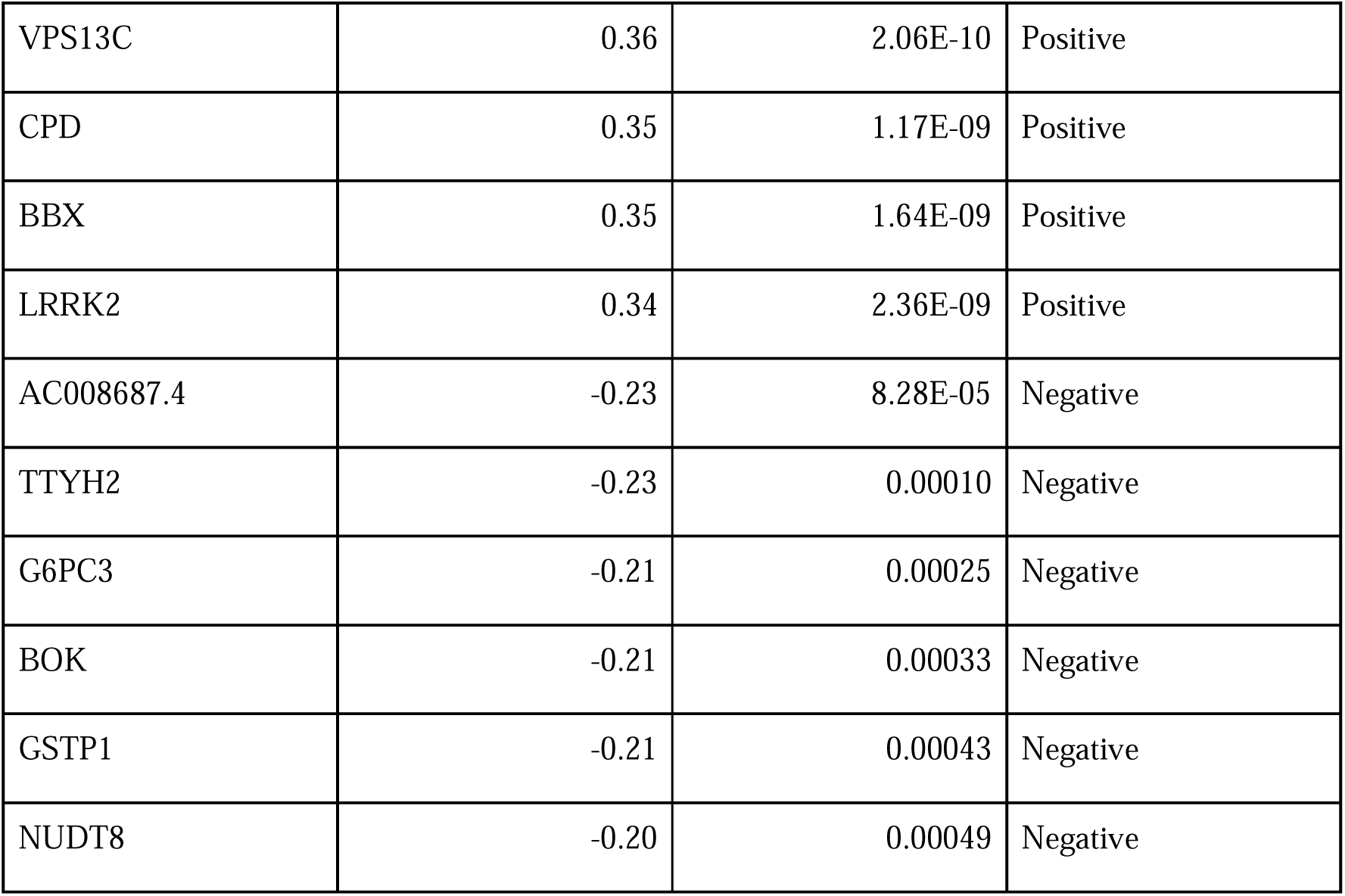
Top Positively and Negatively Correlated Genes Associated with OS Time.

We generated a correlation heatmap to visualize the overall correlation patterns between the identified genes and OS time (**Figure 1a**). This heatmap provides a comprehensive view of the relationships between gene expression levels and OS time. Genes with strong positive correlations are shown in shades of blue, while those with strong negative correlations are represented in shades of red. The intensity of the color reflects the magnitude of the correlation. Afterward, we selected 50 positively correlated genes with high expression levels for enrichment analysis. Significant Gene Ontology (GO) terms included 590 biological processes (BP), 79 cellular components (CC), and no molecular functions (MF). Notable BP enrichments included ‘lipid translocation’ (p = 0.00002) and ‘phospholipid translocation’ (p = 0.00006), while CC analysis highlighted ‘mitochondrial outer membrane’ (p = 0.0002) and ‘organelle outer membrane’ (p = 0.0004). Molecular function analysis revealed significant enrichment in ‘cysteine-type endopeptidase inhibitor activity’ (p = 0.001) and ‘protein serine/threonine kinase activity’ (p = 0.001). Additionally, 246 Reactome pathways were identified, with significant enrichment in pathways such as ‘Ion Transport By P-type ATPases’ (p = 0.0003) and ‘Ion Channel Transport’ (p = 0.0009). The top 5 GO (BP) terms and Reactome pathways are shown in **Figure 1(d)**.

**Figure 1:**
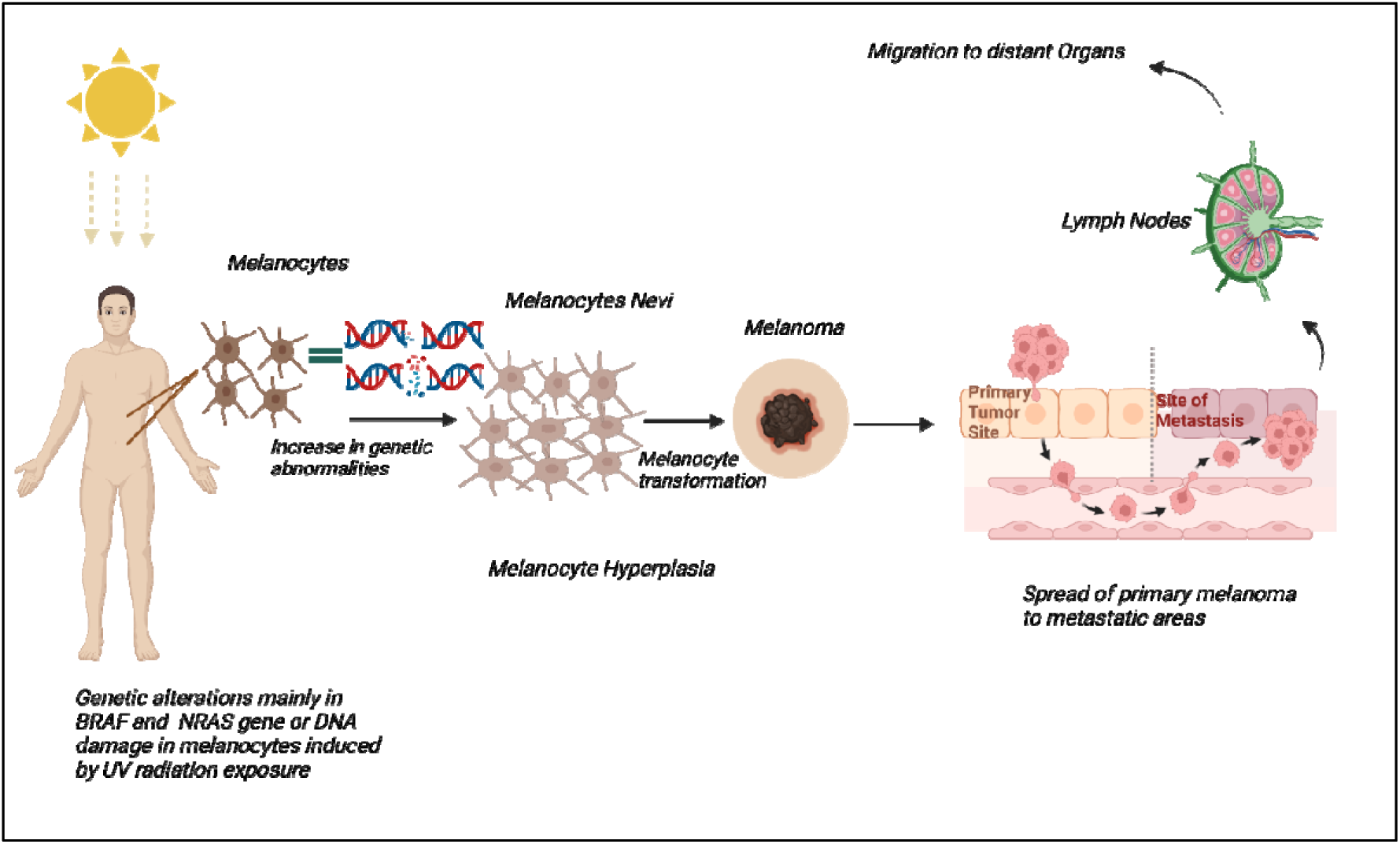
Melanoma development and Progression: The figure depicts the progression from benign to primary melanoma and its spread to metastatic sites

**Figure 2:**
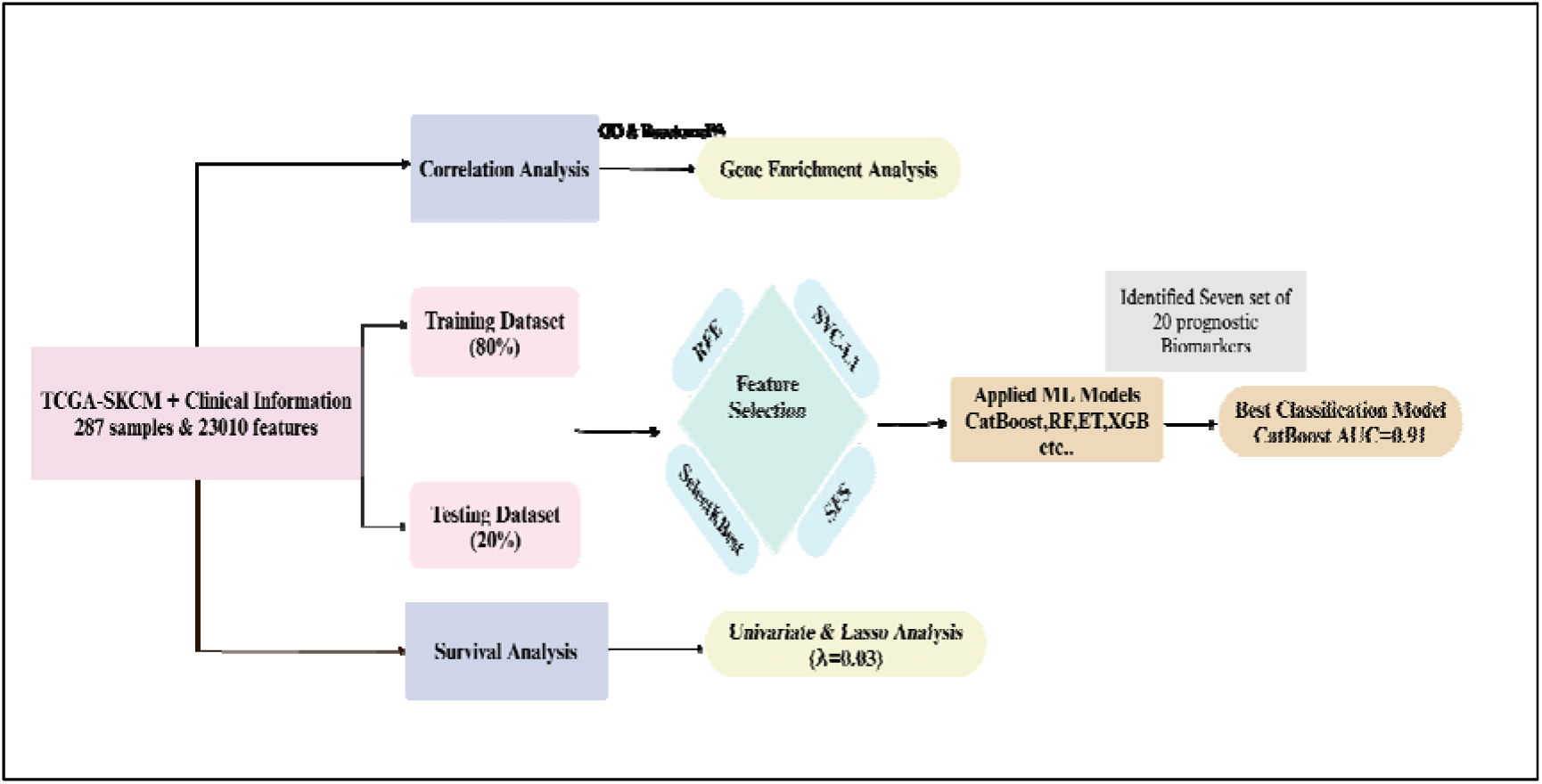
Workflow of our study

**Figure 3.**
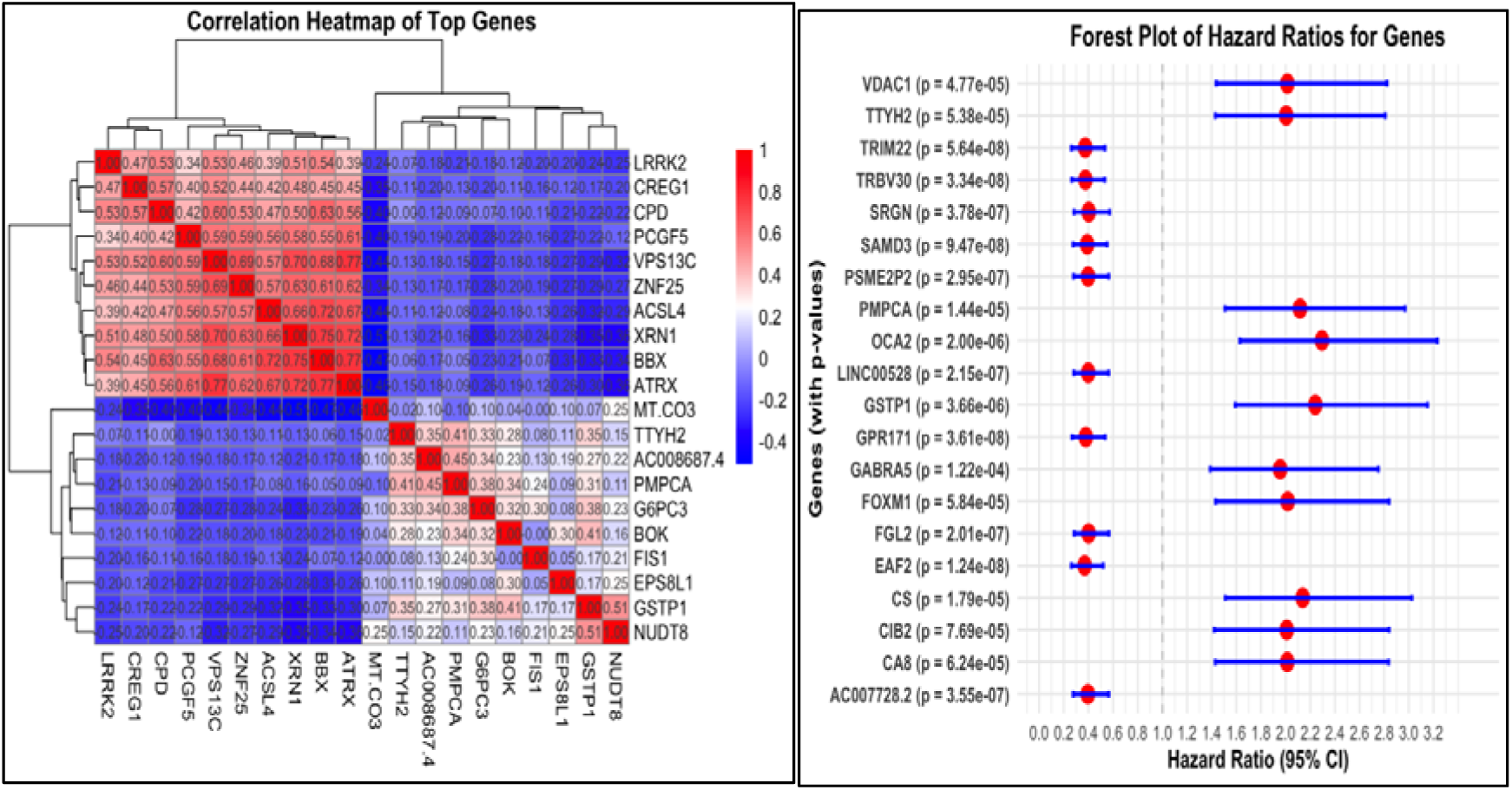
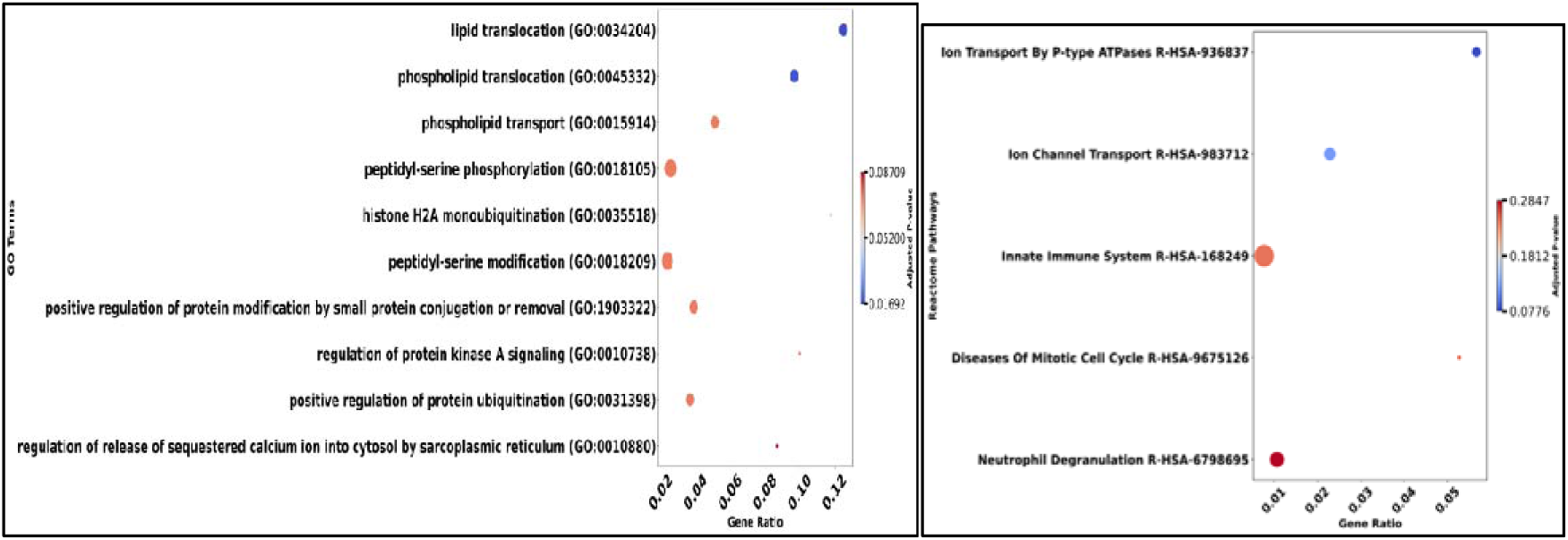
a) The heatmap illustrates the expression patterns of top positive and negative correlated genes b) This forest plot shows the Hazard ratio with a p-value for each gene c) This shows the top 10 Biological processes d) This dot plot shows the top 5 Reactome pathways.

### Survival Analysis

Univariate Cox regression analysis was performed to evaluate the association between gene expression and overall survival. Patients were divided into two groups based on the median OS time, categorizing them into low-risk (G1) and high-risk (G2) groups for each gene. A total of 4,324 genes were identified as statistically significant (p-value < 0.01). Among these, 1,264 genes had a hazard ratio (HR) greater than 1, indicating a potential risk, while 3,060 genes had an HR less than 1, suggesting a protective effect. The top genes with HR values are highlighted in **Figure 1(b)**, while a detailed summary of these findings is presented in **Table 2**.

**Table 2.**
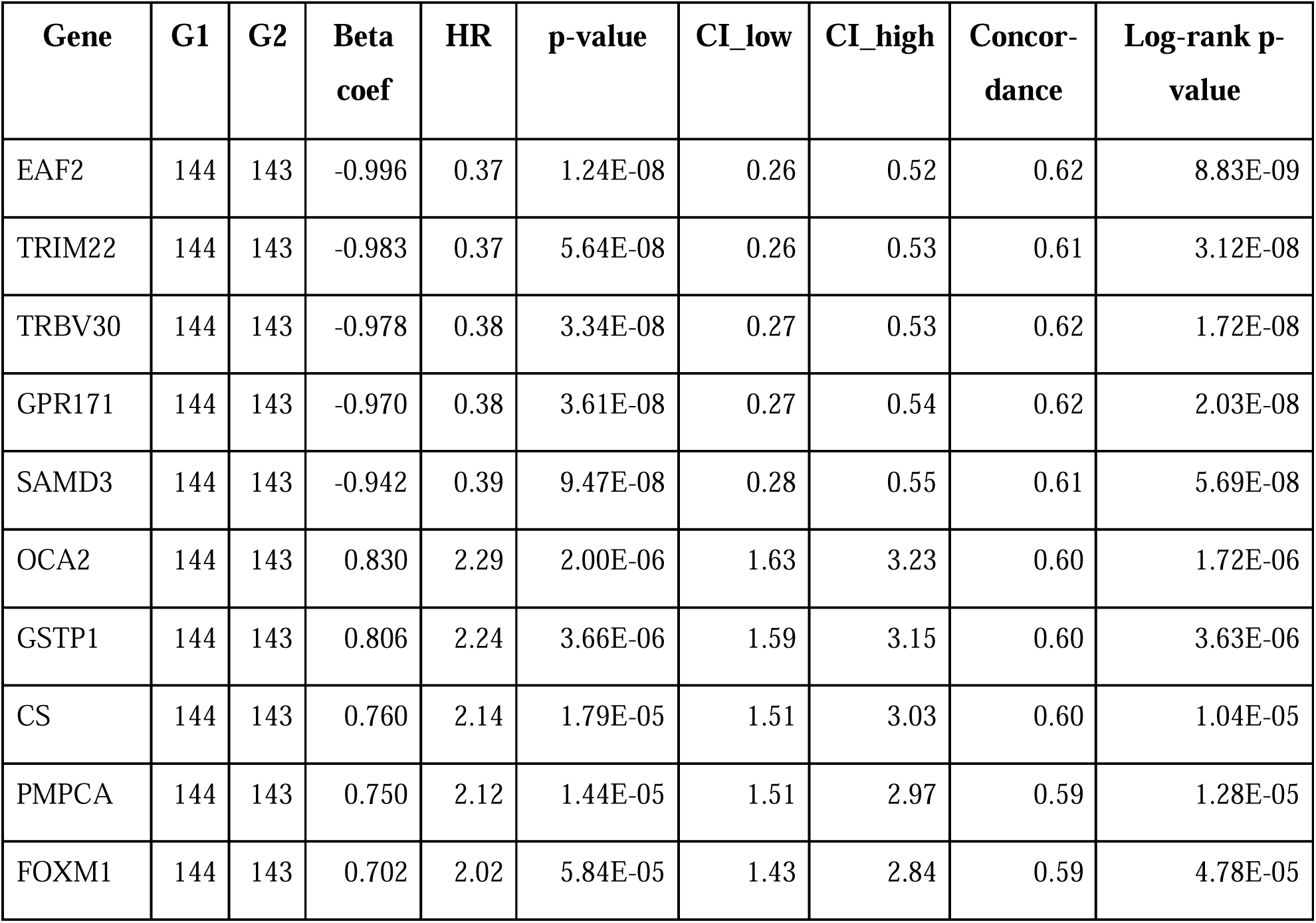
Hazard Ratio (HR)-based Genes Associated with Overall Survival Time.

Subsequently, LASSO Cox regression analysis was performed to identify the most predictive genes for survival outcomes. Using cross-validation, the optimal lambda value was determined to be 0.03, which minimized the cross-validation error. This analysis retained 17 genes (ATP11A, B2M, BISPR, CIB2, CYTL2, GBP1P1, GBP2, GCA, HEXD, HLA.DQB1, KLRC1, LRRK2.DT, MCOLN2, SLC2A5, TTYH2, WIPF1, XCL2) with non-zero coefficients, indicating their significant contribution to OS prediction. Among these, 4 genes were identified as risk factors, and 13 genes were identified as beneficial factors shown in **Table 3**. Additionally, we identified 17 drugs targeting 5 selected genes, detailed in **Supplementary Table S9**.

**Table 3.**
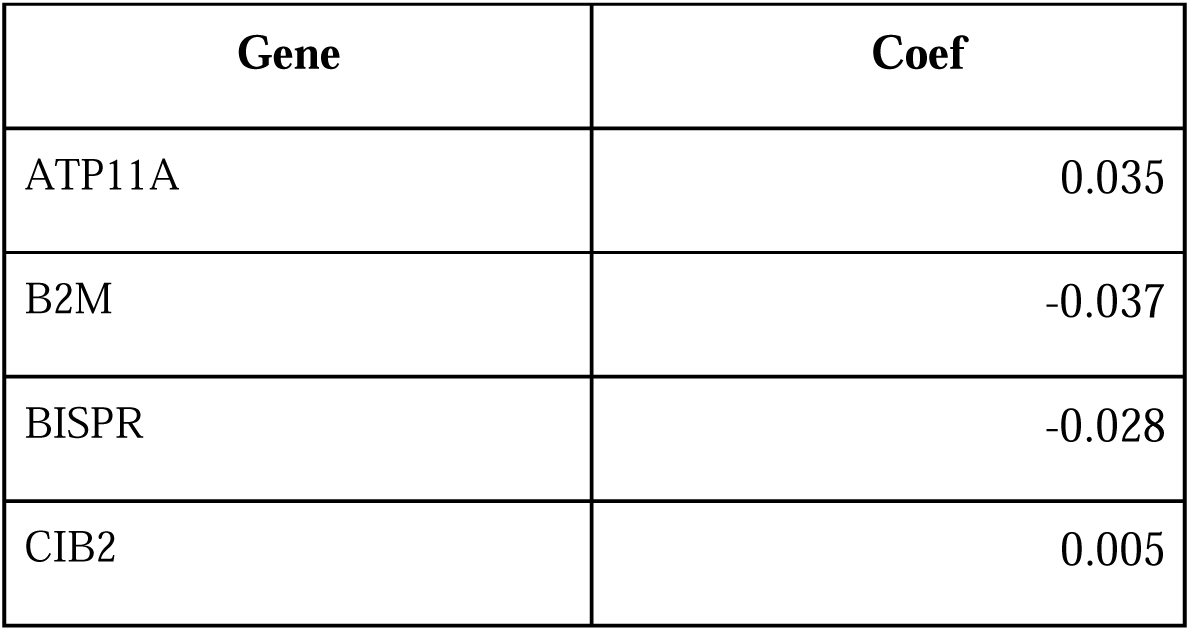
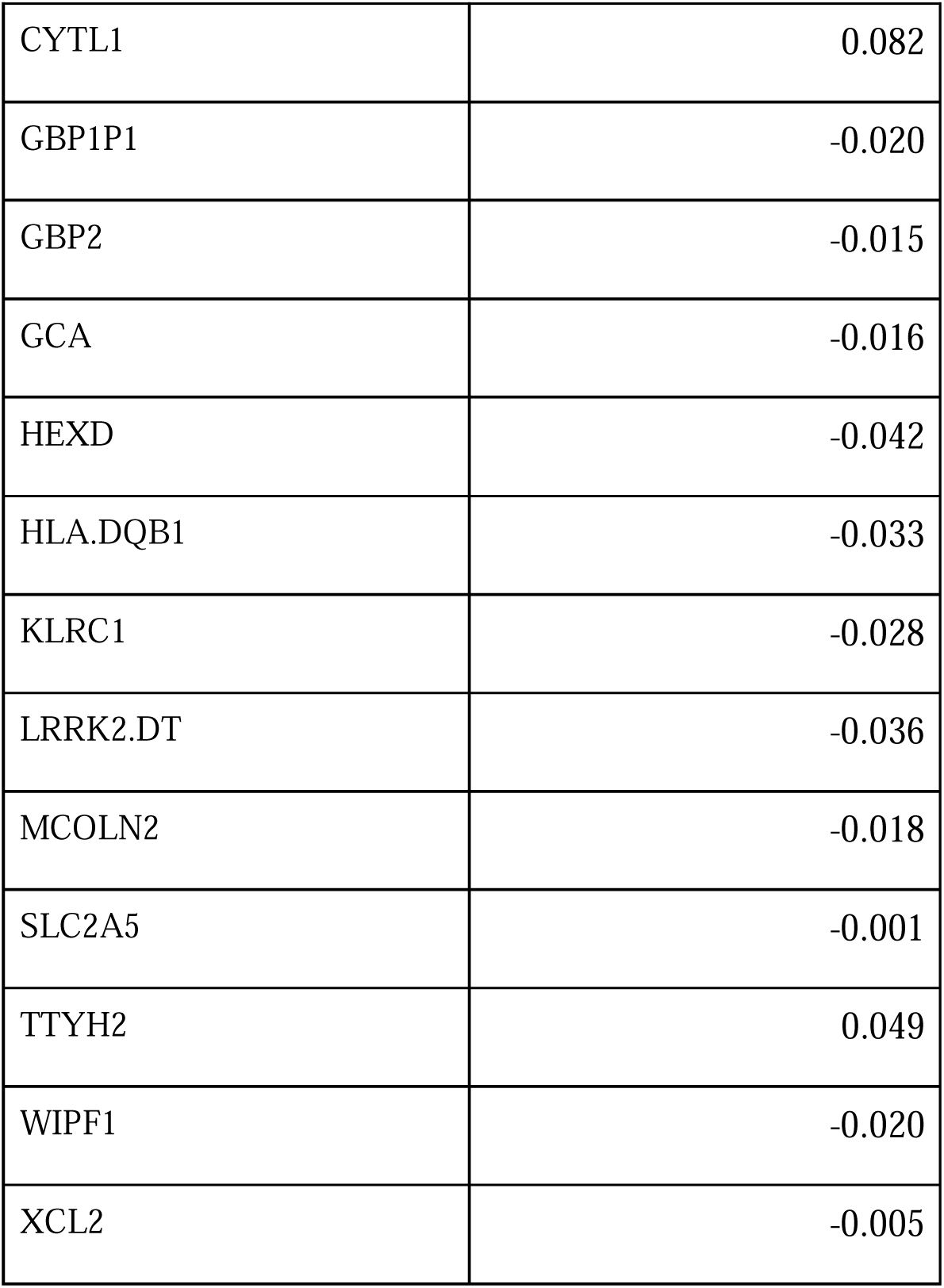
This Table contains 17 Prognosis-related genes in Skin Cutaneous Melanoma.

| Gene | Coef |
| --- | --- |
| ATP11A | 0.035 |
| B2M | -0.037 |
| BISPR | -0.028 |
| CIB2 | 0.005 |
| CYTL1 | 0.082 |
| GBP1P1 | -0.020 |
| GBP2 | -0.015 |
| GCA | -0.016 |
| HEXD | -0.042 |
| HLA.DQB1 | -0.033 |
| KLRC1 | -0.028 |
| LRRK2.DT | -0.036 |
| MCOLN2 | -0.018 |
| SLC2A5 | -0.001 |
| TTYH2 | 0.049 |
| WIPF1 | -0.020 |
| XCL2 | -0.005 |

### Machine Learning Models

To mine important genomic and epigenomic features, we used well-established feature selection methods like SVC-L1(Guyon et al., 2002). SelecKBest (Pedregosa, F. and Varoquaux, G. and Gramfort, A. and Michel et al., 2011) RFE (Panda & Priyadarshi, 2022) and SFS (Esener et al., 2015). Subsequently, prediction models have been developed implementing several machine learning techniques like ExtraTrees (Geurts et al., 2006), CatBoost (Geeitha et al., 2024)

Various feature selection methods were initially applied to identify biomarkers from multiple sets, as detailed in the **Supplementary Table (S2-S8)**. The selected biomarkers were then used to train and evaluate multiple machine-learning models. Performance was assessed using metrics such as accuracy, area under the curve (AUC), specificity, sensitivity, and Matthew’s correlation coefficient (MCC). Among the models, the CatBoost algorithm emerged as the best-performing approach. It achieved an AUC of 0.90 and an MCC of 0.58 on 20 features selected by SVC-L1 based on feature importance, as shown in **Table 4**.

**Table 4.** Performance measures of the 20 mRNA genes selected through SVC-L1 for classification of survival categories in train and test datasets.

| For the primary set of Biomarkers |  |  |  |  |  |  |  |  |  |  |  |  |
| --- | --- | --- | --- | --- | --- | --- | --- | --- | --- | --- | --- | --- |
|  | Train |  |  |  |  |  | Test |  |  |  |  |  |
| Models | Acc | AUC | Sens | Spec | Kappa | MCC | Acc | AUC | Sens | Spec | Kappa | MCC |
| RF | 0.67 | 0.88 | 0.67 | 0.67 | 0.57 | 0.56 | 0.66 | 0.86 | 0.66 | 0.65 | 0.55 | 0.55 |
| SVM | 0.66 | 0.86 | 0.66 | 0.67 | 0.56 | 0.55 | 0.65 | 0.85 | 0.65 | 0.64 | 0.54 | 0.53 |
| ET | 0.73 | 0.91 | 0.73 | 0.73 | 0.65 | 0.64 | 0.7 | 0.87 | 0.7 | 0.69 | 0.61 | 0.61 |
| XGB | 0.66 | 0.86 | 0.66 | 0.67 | 0.55 | 0.55 | 0.66 | 0.84 | 0.66 | 0.65 | 0.55 | 0.55 |
| KNN | 0.6 | 0.81 | 0.6 | 0.61 | 0.48 | 0.47 | 0.57 | 0.79 | 0.57 | 0.58 | 0.44 | 0.42 |
| LightGBM | 0.68 | 0.88 | 0.68 | 0.68 | 0.58 | 0.58 | 0.67 | 0.86 | 0.67 | 0.66 | 0.56 | 0.56 |
| GB | 0.58 | 0.78 | 0.58 | 0.58 | 0.44 | 0.44 | 0.58 | 0.78 | 0.58 | 0.57 | 0.44 | 0.44 |
| AdaBoost | 0.59 | 0.79 | 0.59 | 0.61 | 0.45 | 0.45 | 0.66 | 0.79 | 0.66 | 0.67 | 0.55 | 0.55 |
| **CatBoost** | **0.65** | **0.89** | **0.65** | **0.65** | **0.54** | **0.53** | **0.68** | **0.90** | **0.68** | **0.68** | **0.58** | **0.58** |

For the secondary biomarker set, the CatBoost model achieved an AUC of 0.89, as detailed in **Table 5**. Similarly, for the third biomarker set, CatBoost demonstrated an AUC of 0.87. Consistent performance was observed across the fourth (AUC: 0.85), sixth (AUC: 0.84), and seventh (AUC: 0.89) biomarker sets. Notably, the highest performance was achieved with the fifth biomarker set, where CatBoost attained an AUC of 0.91 and an MCC of 0.64, underscoring its superior predictive capability with this selection of biomarkers.

**Table 5.** Performance measures for a distinct set of biomarkers selected through SVC-L1.

| For the Secondary sets of biomarkers |  |  |  |  |  |  |  |  |  |  |  |  |
| --- | --- | --- | --- | --- | --- | --- | --- | --- | --- | --- | --- | --- |
|  | Train |  |  |  |  |  | Test |  |  |  |  |  |
| Models | Acc | AUC | Sens | Spec | Kappa | MCC | Acc | AUC | Sens | Spec | Kappa | MCC |
| RF | 0.67 | 0.87 | 0.67 | 0.67 | 0.56 | 0.56 | 0.61 | 0.86 | 0.61 | 0.61 | 0.49 | 0.48 |
| SVM | 0.70 | 0.87 | 0.70 | 0.69 | 0.60 | 0.60 | 0.61 | 0.85 | 0.61 | 0.59 | 0.49 | 0.48 |
| ET | 0.75 | 0.91 | 0.75 | 0.76 | 0.67 | 0.67 | 0.67 | 0.89 | 0.67 | 0.66 | 0.56 | 0.56 |
| XGB | 0.66 | 0.85 | 0.66 | 0.66 | 0.55 | 0.54 | 0.61 | 0.84 | 0.61 | 0.60 | 0.49 | 0.48 |
| KNN | 0.59 | 0.82 | 0.59 | 0.60 | 0.47 | 0.46 | 0.51 | 0.78 | 0.51 | 0.49 | 0.36 | 0.35 |
| LightGBM | 0.69 | 0.88 | 0.69 | 0.69 | 0.59 | 0.58 | 0.66 | 0.85 | 0.66 | 0.65 | 0.55 | 0.55 |
| GB | 0.57 | 0.78 | 0.57 | 0.57 | 0.42 | 0.42 | 0.57 | 0.79 | 0.57 | 0.56 | 0.43 | 0.42 |
| MLP | 0.69 | 0.86 | 0.69 | 0.68 | 0.59 | 0.58 | 0.64 | 0.84 | 0.64 | 0.62 | 0.52 | 0.52 |
| AdaBoost | 0.59 | 0.79 | 0.59 | 0.61 | 0.46 | 0.45 | 0.60 | 0.76 | 0.60 | 0.60 | 0.47 | 0.47 |
| **CatBoost** | **0.66** | **0.89** | **0.66** | **0.65** | **0.56** | **0.55** | **0.68** | **0.89** | **0.68** | **0.66** | **0.58** | **0.58** |

| For the third set of Biomarkers |  |  |  |  |  |  |  |  |  |  |  |  |
| --- | --- | --- | --- | --- | --- | --- | --- | --- | --- | --- | --- | --- |
|  | Train |  |  |  |  |  | Test |  |  |  |  |  |
| Models | Acc | AUC | Sens | Spec | Kappa | MCC | Acc | AUC | Sens | Spec | Kappa | MCC |
| RF | 0.66 | 0.87 | 0.66 | 0.67 | 0.56 | 0.55 | 0.64 | 0.84 | 0.64 | 0.64 | 0.52 | 0.52 |
| SVM | 0.65 | 0.85 | 0.65 | 0.65 | 0.54 | 0.53 | 0.63 | 0.84 | 0.63 | 0.61 | 0.51 | 0.50 |
| ET | 0.72 | 0.91 | 0.72 | 0.73 | 0.62 | 0.62 | 0.66 | 0.87 | 0.66 | 0.64 | 0.55 | 0.55 |
| XGB | 0.61 | 0.84 | 0.61 | 0.60 | 0.48 | 0.48 | 0.61 | 0.82 | 0.61 | 0.62 | 0.50 | 0.48 |
| KNN | 0.57 | 0.81 | 0.57 | 0.6 | 0.44 | 0.42 | 0.48 | 0.73 | 0.48 | 0.48 | 0.33 | 0.30 |
| LightGBM | 0.63 | 0.86 | 0.63 | 0.63 | 0.51 | 0.51 | 0.64 | 0.85 | 0.64 | 0.63 | 0.52 | 0.52 |
| GB | 0.52 | 0.75 | 0.52 | 0.53 | 0.37 | 0.36 | 0.6 | 0.77 | 0.6 | 0.60 | 0.47 | 0.47 |
| MLP | 0.67 | 0.83 | 0.67 | 0.67 | 0.57 | 0.56 | 0.63 | 0.8 | 0.63 | 0.60 | 0.51 | 0.50 |
| AdaBoost | 0.54 | 0.78 | 0.54 | 0.58 | 0.40 | 0.39 | 0.58 | 0.77 | 0.58 | 0.63 | 0.46 | 0.44 |
| **CatBoost** | **0.66** | **0.89** | **0.66** | **0.67** | **0.56** | **0.55** | **0.66** | **0.87** | **0.66** | **0.68** | **0.56** | **0.55** |
| For the fourth set of Biomarkers |  |  |  |  |  |  |  |  |  |  |  |  |
|  | Train |  |  |  |  |  | Test |  |  |  |  |  |
| Models | Acc | AUC | Sens | Spec | Kappa | MCC | Acc | AUC | Sens | Spec | Kappa | MCC |
| RF | 0.66 | 0.86 | 0.66 | 0.66 | 0.55 | 0.55 | 0.53 | 0.82 | 0.53 | 0.53 | 0.38 | 0.38 |
| SVM | 0.66 | 0.84 | 0.66 | 0.66 | 0.55 | 0.54 | 0.60 | 0.79 | 0.60 | 0.60 | 0.47 | 0.47 |
| ET | 0.68 | 0.89 | 0.68 | 0.69 | 0.58 | 0.58 | 0.58 | 0.84 | 0.58 | 0.57 | 0.44 | 0.44 |
| XGB | 0.61 | 0.82 | 0.61 | 0.61 | 0.48 | 0.48 | 0.57 | 0.82 | 0.57 | 0.57 | 0.43 | 0.42 |
| KNN | 0.50 | 0.78 | 0.50 | 0.55 | 0.35 | 0.33 | 0.49 | 0.75 | 0.49 | 0.46 | 0.33 | 0.32 |
| LightGBM | 0.61 | 0.84 | 0.61 | 0.61 | 0.49 | 0.48 | 0.56 | 0.81 | 0.56 | 0.55 | 0.41 | 0.41 |
| GB | 0.54 | 0.77 | 0.54 | 0.54 | 0.39 | 0.39 | 0.52 | 0.76 | 0.52 | 0.51 | 0.37 | 0.36 |
| MLP | 0.69 | 0.85 | 0.69 | 0.69 | 0.59 | 0.58 | 0.63 | 0.79 | 0.63 | 0.62 | 0.50 | 0.50 |
| AdaBoost | 0.54 | 0.75 | 0.54 | 0.54 | 0.39 | 0.38 | 0.42 | 0.71 | 0.42 | 0.41 | 0.23 | 0.23 |
| **CatBoost** | **0.67** | **0.88** | **0.67** | **0.67** | **0.56** | **0.56** | **0.60** | **0.85** | **0.60** | **0.61** | **0.48** | **0.47** |

**For the fifth set of Biomarkers**
|  | Train |  |  |  |  |  | Test |  |  |  |  |  |
| --- | --- | --- | --- | --- | --- | --- | --- | --- | --- | --- | --- | --- |
| Models | Acc | AUC | Sens | Spec | Kappa | MCC | Acc | AUC | Sens | Spec | Kappa | MCC |
| RF | 0.65 | 0.85 | 0.65 | 0.64 | 0.53 | 0.53 | 0.73 | 0.87 | 0.73 | 0.72 | 0.64 | 0.64 |
| SVM | 0.61 | 0.83 | 0.61 | 0.60 | 0.48 | 0.48 | 0.61 | 0.82 | 0.61 | 0.59 | 0.49 | 0.48 |
| ET | 0.71 | 0.90 | 0.71 | 0.72 | 0.62 | 0.62 | 0.69 | 0.89 | 0.69 | 0.68 | 0.59 | 0.59 |
| XGB | 0.61 | 0.83 | 0.61 | 0.60 | 0.48 | 0.48 | 0.61 | 0.85 | 0.61 | 0.62 | 0.49 | 0.48 |
| KNN | 0.56 | 0.78 | 0.56 | 0.56 | 0.42 | 0.42 | 0.51 | 0.75 | 0.51 | 0.50 | 0.36 | 0.35 |
| LightGBM | 0.64 | 0.85 | 0.64 | 0.64 | 0.53 | 0.52 | 0.64 | 0.86 | 0.64 | 0.63 | 0.52 | 0.52 |
| GB | 0.56 | 0.77 | 0.56 | 0.56 | 0.41 | 0.41 | 0.58 | 0.81 | 0.58 | 0.58 | 0.44 | 0.44 |
| MLP | 0.67 | 0.84 | 0.67 | 0.66 | 0.56 | 0.56 | 0.61 | 0.77 | 0.61 | 0.62 | 0.49 | 0.48 |
| CatBoost | 0.66 | 0.89 | 0.66 | 0.64 | 0.55 | 0.54 | 0.73 | 0.91 | 0.73 | 0.72 | 0.64 | 0.64 |
| For the sixth set of Biomarkers |  |  |  |  |  |  |  |  |  |  |  |  |
|  | Train |  |  |  |  |  | Test |  |  |  |  |  |
| Models | Acc | AUC | Sens | Spec | Kappa | MCC | Acc | AUC | Sens | Spec | Kappa | MCC |
| RF | 0.65 | 0.86 | 0.65 | 0.67 | 0.54 | 0.54 | 0.56 | 0.81 | 0.56 | 0.58 | 0.41 | 0.41 |
| SVM | 0.58 | 0.77 | 0.58 | 0.59 | 0.45 | 0.44 | 0.49 | 0.72 | 0.49 | 0.50 | 0.32 | 0.32 |
| ET | 0.69 | 0.89 | 0.69 | 0.70 | 0.59 | 0.59 | 0.65 | 0.84 | 0.65 | 0.66 | 0.53 | 0.53 |
| XGB | 0.60 | 0.83 | 0.60 | 0.60 | 0.47 | 0.47 | 0.55 | 0.79 | 0.55 | 0.57 | 0.40 | 0.39 |
| KNN | 0.63 | 0.85 | 0.63 | 0.65 | 0.52 | 0.51 | 0.60 | 0.81 | 0.60 | 0.64 | 0.47 | 0.47 |
| LightGBM | 0.56 | 0.80 | 0.56 | 0.55 | 0.41 | 0.41 | 0.61 | 0.79 | 0.61 | 0.61 | 0.49 | 0.48 |
| GB | 0.66 | 0.84 | 0.66 | 0.66 | 0.55 | 0.54 | 0.67 | 0.80 | 0.67 | 0.67 | 0.56 | 0.56 |
| MLP | 0.63 | 0.86 | 0.63 | 0.63 | 0.51 | 0.51 | 0.55 | 0.82 | 0.55 | 0.54 | 0.40 | 0.39 |
| CatBoost | 0.65 | 0.86 | 0.65 | 0.67 | 0.54 | 0.54 | 0.56 | 0.81 | 0.56 | 0.58 | 0.41 | 0.41 |
| For the seventh set of Biomarkers |  |  |  |  |  |  |  |  |  |  |  |  |
| Models | Acc | AUC | Sens | Spec | Kappa | MCC | Acc | AUC | Sens | Spec | Kappa | MCC |
| RF | 0.64 | 0.87 | 0.64 | 0.64 | 0.52 | 0.52 | 0.63 | 0.86 | 0.63 | 0.62 | 0.50 | 0.50 |
| SVM | 0.61 | 0.80 | 0.61 | 0.63 | 0.48 | 0.47 | 0.61 | 0.74 | 0.61 | 0.61 | 0.49 | 0.48 |
| ET | 0.70 | 0.89 | 0.70 | 0.70 | 0.60 | 0.59 | 0.65 | 0.87 | 0.65 | 0.64 | 0.53 | 0.53 |
| XGB | 0.63 | 0.85 | 0.63 | 0.63 | 0.50 | 0.50 | 0.55 | 0.83 | 0.55 | 0.53 | 0.40 | 0.39 |
| KNN | 0.56 | 0.80 | 0.56 | 0.56 | 0.43 | 0.42 | 0.47 | 0.75 | 0.47 | 0.45 | 0.30 | 0.29 |
| LightGBM | 0.66 | 0.86 | 0.66 | 0.66 | 0.54 | 0.54 | 0.59 | 0.83 | 0.59 | 0.59 | 0.46 | 0.45 |
| GB | 0.57 | 0.79 | 0.57 | 0.57 | 0.43 | 0.43 | 0.60 | 0.81 | 0.60 | 0.62 | 0.47 | 0.47 |
| MLP | 0.69 | 0.86 | 0.69 | 0.70 | 0.60 | 0.59 | 0.61 | 0.80 | 0.61 | 0.60 | 0.49 | 0.48 |
| **CatBoost** | **0.65** | **0.89** | **0.65** | **0.64** | **0.54** | **0.53** | **0.64** | **0.89** | **0.64** | **0.63** | **0.52** | **0.52** |

The classification performance of the CatBoost model was initially evaluated on the primary set, where it achieved outstanding results with AUC values of 0.99 for Class 0, 0.83 for Class 1, 0.93 for Class 2, and 0.84 for Class 3. Subsequently, the model was tested on the additional set for seven distinct biomarker sets, with AUC values reported for each class. For the secondary set of biomarkers, the model achieved AUC values of 0.99 for Class 0, 0.77 for Class 1, 0.90 for Class 2, and 0.90 for Class 3. In the third set of biomarkers, the AUC values were 0.98 for Class 0, 0.80 for Class 1, 0.93 for Class 2, and 0.73 for Class 3. The fourth set of biomarkers recorded AUC values of 0.80, 0.85, 0.90, and 0.87 for Classes 0, 1, 2, and 3, respectively. Similarly, the fifth set of biomarkers showed strong performance with AUC values of 0.99 for Class 0, 0.85 for Class 1, 0.91 for Class 2, and 0.90 for Class 3. The sixth set of biomarkers achieved AUC values of 0.92 for Class 0, 0.75 for Class 1, 0.90 for Class 2, and 0.77 for Class 3. Finally, the seventh set of biomarkers demonstrated AUC values of 0.95 for Class 0, 0.82 for Class 1, 0.93 for Class 2, and 0.85 for Class 3. These results, shown in **Table 6**, and the corresponding AUC curve presented in **Figure 5**, highlight the robust performance of the CatBoost model across multiple biomarkers sets, with variations in AUC values reflecting differences in the predictive potential of the biomarker sets.

**Table 6.**
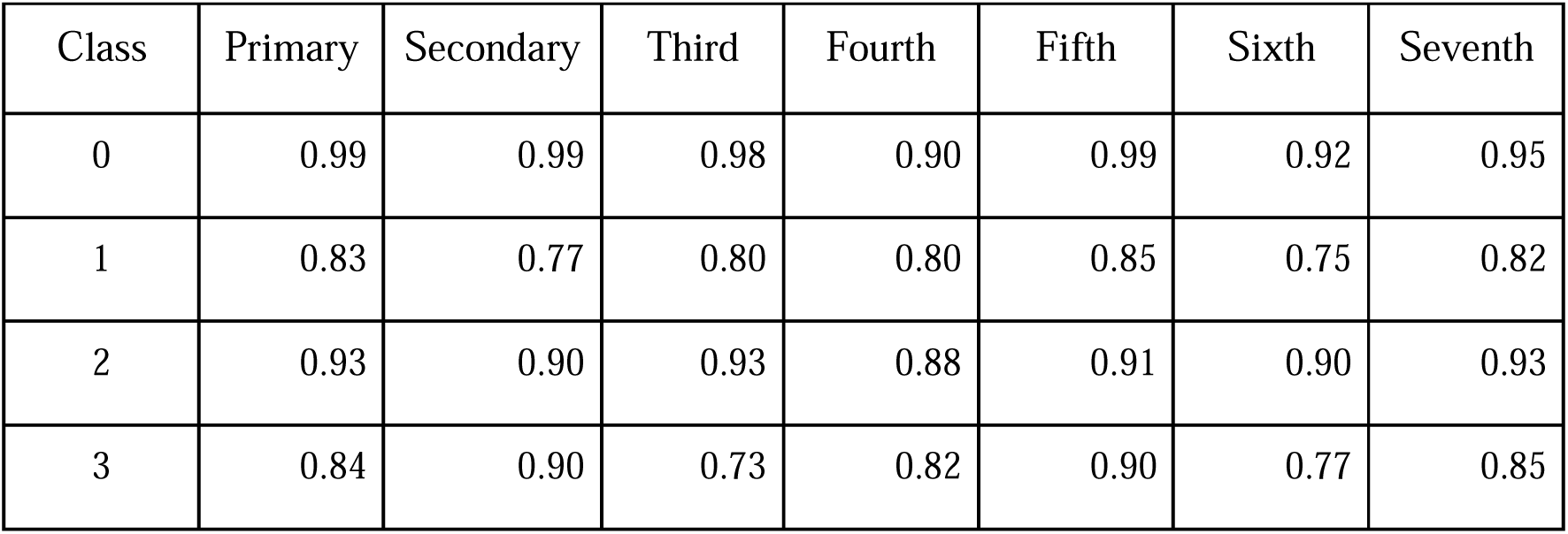
AUC for Each Class Using the Best-Performing Model on All Seven Sets of Biomarkers.

**Figure 4:**
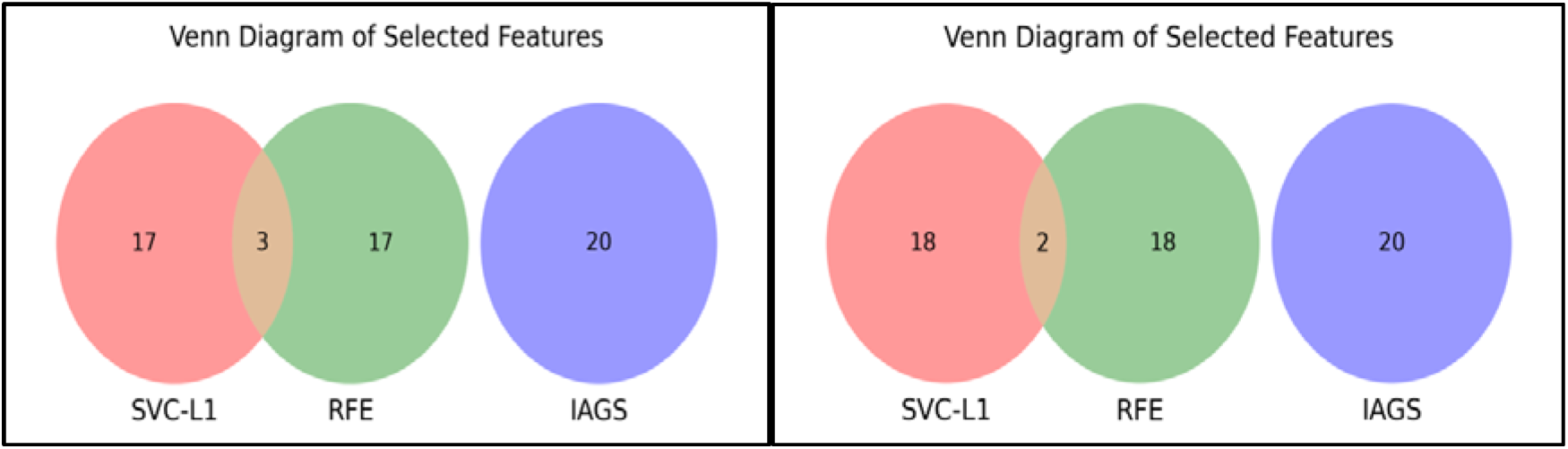
a) Venn diagram showing the common features selected by SVC-L1 and RFE in the primary set of biomarkers. b) Venn diagram depicting the common features selected by SVC-L1 and RFE in the second set of biomarkers.

**Figure 5.**
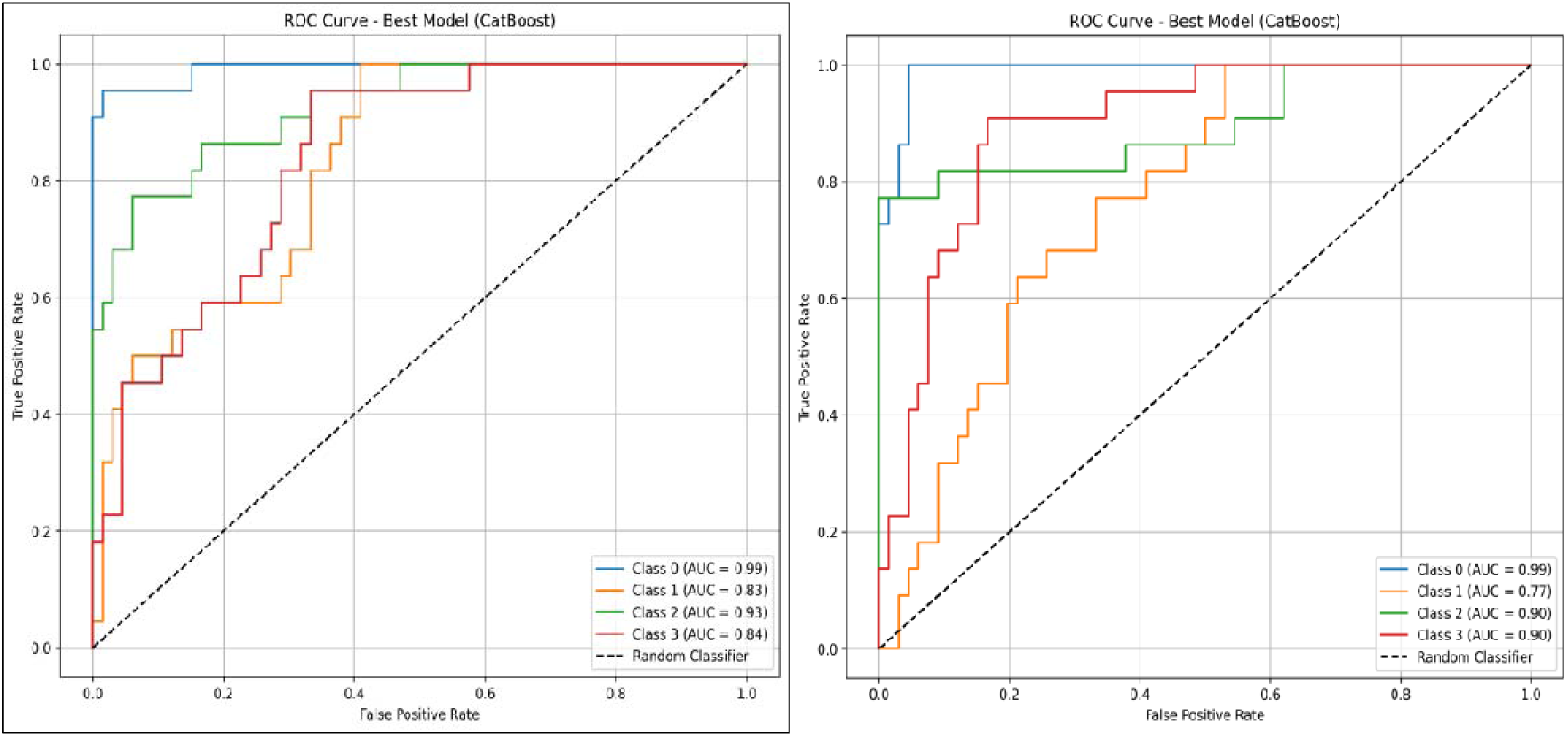
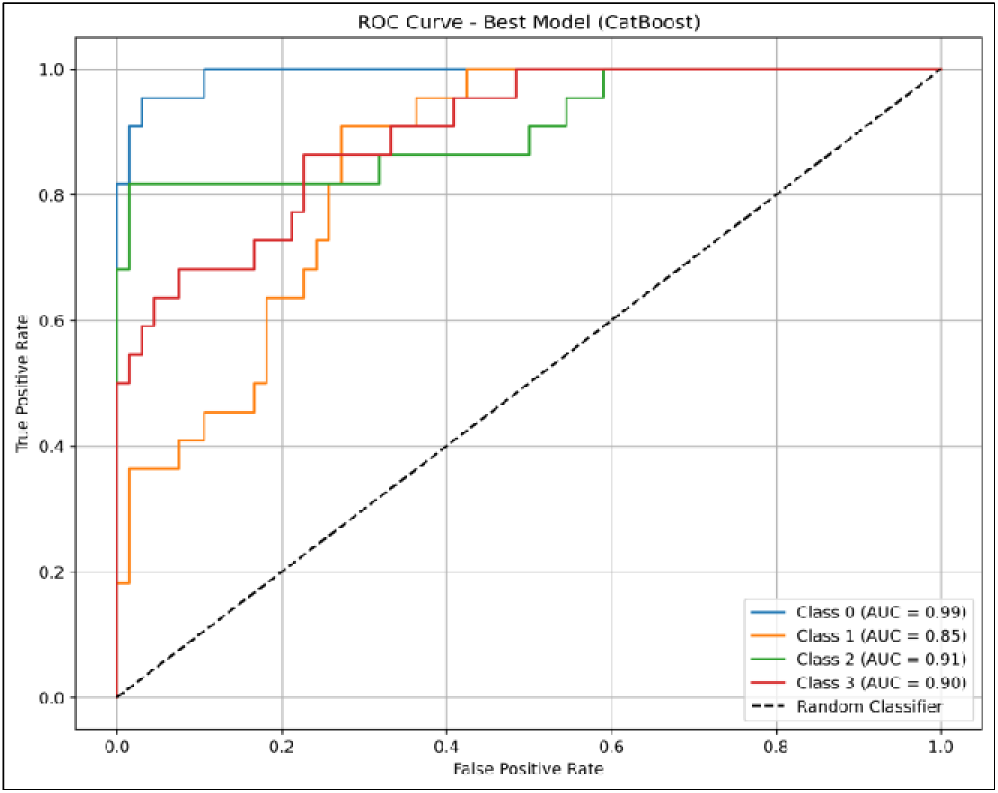
AUC-ROC curve demonstrating the classification performance of the CatBoost model for each survival group: a) For the primary set of biomarkers, the AUC curve classifies overall survival into four categories. b) For the secondary set of biomarkers, the AUC curve shows classification performance for each survival class. c) For the fifth set of biomarkers, the AUC curve classifies survival time into Class (0-3)

The AUC values in **Table 6** illustrate the predictive performance of the best-performing model across seven biomarker sets for survival classes (0–1 year, 1–3 years, 3–5 years, and >5 years). For Class 0 (0–1 year survival), the AUC values are consistently high, ranging from 0.90 to 0.99, with the primary, secondary, and fifth biomarker sets achieving the highest AUC (0.99). This indicates exceptional accuracy in identifying patients with the shortest survival times, highlighting these biomarker sets as highly effective for predicting high-risk patients who may require immediate clinical intervention. For Class 1 (1–3 years survival), the AUC values are moderate, ranging from 0.75 to 0.85, with the fifth biomarker set demonstrating the highest predictive accuracy (0.85) and the sixth set performing the least effectively (0.75). These results suggest moderate success in predicting mid-range survival groups, indicating room for improvement in this category.

For Class 2 (3–5 years survival), the AUC values are consistently high across all biomarkers sets, ranging from 0.88 to 0.93, with the primary, third, and seventh sets achieving the highest AUC (0.93). This consistency demonstrates the robustness of these sets in predicting intermediate survival times, making them promise for accurate prognostic stratification in this group. For Class 3 (>5 years survival), the AUC values show more variation, ranging from 0.73 to 0.90. The secondary and fifth biomarker sets achieve the highest performance (0.90), while the third set shows the lowest (0.73), indicating that while some sets are effective for long-term survival prediction, others may lack the necessary features to differentiate this class accurately.

Overall, the primary biomarker set stands out for its exceptional performance across all classes, particularly for Class 0 and Class 2 (AUC = 0.99 and 0.93, respectively), making it the most reliable set for survival prediction. The fifth biomarker set demonstrates high performance, particularly for Classes 0 and 3 (AUC = 0.99 and 0.90), showing its potential for identifying high-risk and long-term survivors. The results reveal that the highest predictive accuracy is observed for Class 0, suggesting the model excels in identifying high-risk patients. Moderate performance for Class 1 indicates challenges in predicting mid-range survival groups, while high accuracy for Class 2 and variable performance for Class 3 highlights the model’s strengths. These findings emphasize the potential of these biomarker sets in predicting survival outcomes in SKCM patients, with the primary and fifth sets showing the best performance (Fig 5)

### Ensemble Models

The Voting and Stacking classifiers were applied to the 20 prognostic genes from the primary set. The Voting classifier combined RF, ET, and LightGBM, achieved an AUC of 0.87 and an MCC of 0.56. The Stacking classifier was evaluated using Logistic Regression as the meta-model with RF and ET as base models and CatBoost as the meta-model with RF, ET, and XGB as base models. These stacking models achieved AUC values of 0.88, with MCC scores of 0.61. The results are summarized in **Table 7**.

**Table 7.** Performance on 20 genes selected using SVC-L1.

| Combination of RF, ET & LightGBM |  |  |  |  |  |  |  |  |  |  |  |  |
| --- | --- | --- | --- | --- | --- | --- | --- | --- | --- | --- | --- | --- |
|  | Train |  |  |  |  |  | Test |  |  |  |  |  |
| Classifier | Acc | AUC | Sens | Spec | Kappa | MCC | Acc | AUC | Sens | Spec | Kappa | MCC |
| Voting | 0.69 | 0.90 | 0.69 | 0.70 | 0.59 | 0.59 | 0.67 | 0.87 | 0.67 | 0.66 | 0.56 | 0.56 |

| LR as meta-model & RF, ET as base-models |  |  |  |  |  |  |  |  |  |  |  |  |
| --- | --- | --- | --- | --- | --- | --- | --- | --- | --- | --- | --- | --- |
| Stacking | 0.74 | 0.91 | 0.74 | 0.74 | 0.66 | 0.65 | 0.70 | 0.88 | 0.70 | 0.70 | 0.61 | 0.61 |
| CatBoost as meta-model & RF, ET, & XGB as base-models |  |  |  |  |  |  |  |  |  |  |  |  |
| Stacking | 0.69 | 0.89 | 0.69 | 0.71 | 0.59 | 0.59 | 0.64 | 0.88 | 0.64 | 0.64 | 0.52 | 0.52 |

The best existing method, which focused on IAGS, achieved an AUC of 0.88. In our study, four out of seven biomarker sets performed even better, with AUC values higher than 0.88. This demonstrates that our approach is more effective in identifying prognostic biomarkers with higher accuracy.

## Discussion

Melanoma is the most lethal and deadliest form of skin cancer (Lombard et al., 2019). The prognosis is often poor due to early metastasis, which remains the primary cause of death in affected individuals (Sun et al., 2019). Therefore, early detection of SKCM and effective stratification of risk assessment are crucial for timely treatment and the improvement of survival rates. As one of the most immunogenic tumors, the role of immune regulation and the potential of immunotherapy in SKCM have consistently been central to research and clinical discussions (Marzagalli et al., 2019). We utilized the TCGA dataset to identify potential prognostic biomarkers for SKCM. Through in silico analyses of gene expression profiles from TCGA, we aimed to uncover key genes associated with patient survival, disease progression, and therapeutic response, ultimately enhancing the prognostic accuracy for SKCM patients.

Our study takes a comprehensive approach, focusing on a broad range of genes rather than targeting specific types or pathways. This strategy aims to identify distinct sets of biomarkers associated with overall survival in SKCM. We analyzed all available genes using statistical and machine-learning methods to uncover meaningful prognostic biomarkers. Based on the median OS time, a cohort of 287 patients with available survival data was divided into high- and low-risk groups. In our analysis, we first identified genes correlated with OS time. Pearson correlation analysis revealed several genes with significant correlations to OS, including CREG1 and PCGF5, which showed positive correlations, and AC008687.4 and TTYH2, which demonstrated negative correlations. Next, we selected the top 50 positively correlated genes for further exploration. These genes were enriched in various biological processes and pathways, including lipid translocation and phospholipid translocation. Reactome pathway analysis further revealed their involvement in key pathways such as *’*Ion Transport By P-type ATPases’ and ‘Ion Channel Transport.’

Our univariate Cox regression analysis revealed a substantial number of genes significantly associated with OS time, with a total of 4324 genes found to have p-values below 0.01. Among these, we identified 1264 genes associated with a hazard ratio (HR>1), indicating them as potential risk factors, while 3060 genes were associated with HR<1, suggesting a protective effect on survival. These results emphasize the complexity of SKCM’s molecular landscape, where multiple genes may promote or inhibit tumor progression and patient survival. We further refined our gene list to 17 genes with non-zero coefficients by applying LASSO Cox regression. The identification of risk and beneficial factors among the 17 selected genes is another key finding of our study. Four of these genes were classified as risk factors, while the remaining 13 genes were classified as beneficial factors. This classification not only underscores the diverse nature of the molecular drivers of SKCM but also offers potential biomarkers for therapeutic targeting. For instance, CYTL1 expression was progressively upregulated in normal skin, nevi or malignant nevi, and melanoma (Tao et al., 2023). Similarly, B2M (Beta-2-Microglobulin) is associated with worse OS and disease-specific survival (DSS) in SKCM, acting as a hazardous factor by contributing to tumor progression and immune evasion (H. Zhang et al., 2021) GBP2 was down-regulated in SKCM and Low GBP2 expression was positively correlated with poor prognosis of SKCM: (Ji et al., 2021). The Expressions of HLA Class II (HLA-DQB1) Genes Were Up-Regulated in Cutaneous Melanoma (Chen et al., 2019). KLRC1 expression in SKCM is associated with immune cell infiltration, highlighting its role in immune surveillance and tumor microenvironment modulation (Kamiya et al., 2019) MCOLN2 (TRPML2) has not been extensively studied in SKCM. However, its upregulation in other cancers like prostate tumors and its association with poor prognosis suggests it might play a role in melanoma progression by potentially influencing tumor growth, invasion, or immune evasion (li & Zhu, 2023) in three distinct malignancies that contained SKCM, SARC, and THCA, SLC2A5 acted as a protective factor (Liu et al., 2024). WIPF1 up regulated in early melanoma (Zia et al., 2023). XCL2 shows high expression in SKCM (Xiong et al., 2020). To further enhance the clinical applicability of our findings, we also explored drug-target prediction using the DGIdb database. We identified 17 drugs targeting 5 of the most relevant genes. This step underscores the clinical relevance of our biomarker discovery process, suggesting that the genes we identified are predictive of survival outcomes and potentially actionable through existing or future therapeutic agents. Incorporating drug-target information into our model provides an additional layer of clinical utility, enabling the identification of drugs that may benefit patients based on their gene expression profiles. The drugs identified in this study may offer opportunities for personalized treatment strategies, particularly for those with high-risk profiles based on the gene expression signature.

The top 20 mRNA genes selected using SVC-L1 achieved an AUC of 0.91, indicating that machine-learning techniques can be effectively utilized for survival prediction in SKCM. These results are consistent with other machine learning-based studies in melanoma, which have demonstrated the utility of advanced algorithms, such as support vector machines (SVM) and random forests, for identifying prognostic biomarkers (Bhalla et al., 2019). So, the primary set of biomarkers associated with skin cutaneous melanoma includes TEKT5, ZNF154, H2AC14, BX284668.6, MYCNOS, STUM, SERTM2, RPSAP18, REG4, PSCA, PAEP, ACTR3C, MSLN, MRPS18AP1, ISLR, IL37, IGLV3.16, H2BC11, GPR25, and MTND4P35. PSCA (Prostate Stem Cell Antigen) is upregulated in SKCM, as suggested by its positive correlation with genes related to RNA modifications, indicating an active role in the tumor’s biological processes (C. Wang et al., 2024). PAEP (Progestogen-Associated Endometrial Protein) is upregulated and correlates with poor survival outcomes in SKCM (J. Yan et al., 2020). ACTR3C mutations, such as p. Gly58Arg or p.Gly58Glu, might alter protein function and affect key cellular processes such as cell signaling, motility, or adhesion, which are crucial in melanoma progression (Srinivasan et al., 2021). MSLN expression is downregulated in SKCM (Li et al., 2022). ISLR is also downregulated and negatively correlated with Tumor Mutation load (TMB) (C. Zhang et al., 2021). STUM is a biomarker for response to treatment with Nivolumab (immune checkpoint inhibitors) (Bannon et al., 2024). SERTM2 gene alterations are frequently observed in melanoma, suggesting a potential role in tumor progression and immune evasion. These alterations may impact pathways related to melanoma growth, metastasis, and immune response modulation (Niu et al., 2024). Secondary biomarkers play a vital role in other cancers but are not directly related to to SKCM (Bisaccia et al., 2023; J. He et al., 2020; Jia et al., 2021; Larsson et al., 2022; Severgnini et al., 2022). For the fifth set of biomarkers, we did not find any direct evidence of their specific role in SKCM. These genes are newly identified in our study and may serve as potential prognostic biomarkers (S. Ding et al., 2022; Fang et al., 2022; JING HUI NANA SUN, 2024; W. Yang et al., 2021). For Class 0 (0–1 year survival), the AUC values were exceptionally high, ranging from 0.90 to 0.99, with the primary, secondary, and fifth biomarker sets achieving the highest AUC of 0.99. This suggests that these sets are highly effective in identifying high-risk patients with the shortest survival times, making them valuable for early-stage risk assessment and timely clinical intervention. In contrast, for Class 1 (1–3 years survival), the AUC values ranged from 0.75 to 0.85, with the fifth biomarker set showing the best performance (AUC = 0.85). The moderate AUC for this class indicates that predicting mid-range survival times remains challenging, and further refinement of the biomarkers may be needed to improve accuracy in this group.

For Class 2 (3–5 years survival), the AUC values were consistently high across all sets, ranging from 0.88 to 0.93, with the primary, third, and seventh sets achieving the highest AUC of 0.93. This highlights the robustness of these biomarker sets in predicting intermediate survival, making them useful for more accurate prognostic stratification. However, for Class 3 (>5 years survival), the AUC values showed more variation, ranging from 0.73 to 0.90, with the secondary and fifth sets performing the best (AUC = 0.90). The lower performance for some sets in this class suggests that further refinement is needed to capture the specific molecular features associated with long-term survival. Overall, the primary and fifth biomarker sets demonstrated superior performance, particularly for high-risk and long-term survival predictions, showing promise for clinical integration to enhance patient management strategies in SKCM.

A key limitation of this study is that it is purely bioinformatics in nature, relying on computational analysis of existing datasets. While the results are promising, they should be considered as preliminary findings. Further experimental validation in wet lab settings, including gene expression profiling in patient samples, is crucial to confirm the biological relevance of the identified biomarkers. This would provide stronger evidence for their clinical applicability and utility in the prognosis and treatment of SKCM.

## Conclusion

This study identified seven distinct sets of biomarkers and developed a robust prognostic model for predicting overall survival in skin cutaneous melanoma. By utilizing machine learning techniques, we were able to select the most relevant biomarkers that contribute to patient prognosis. The model showed promising results in classifying patients into different survival categories based on OS time, thereby offering significant value for clinical prognosis prediction. Furthermore, the findings highlight the potential of these biomarkers for guiding personalized treatment strategies, improving patient outcomes, and informing therapeutic decisions in SKCM. This approach demonstrates the possibility of identifying different sets of biomarkers, each contributing to the prognostic landscape of SKCM. However, further validation and clinical trials are needed to confirm the clinical applicability of these biomarkers in real-world settings.

## Abbreviations

SKCM: Skin Cutaneous Melanoma
OS: Overall survival
IAGS: Invasion-associated genes
TCGA: The Cancer Genome Atlas
FRG: Ferroptosis-related gene
DGIdb: Drug Gene Interaction Database
ML: Machine Learning
AUROC: Area Under the Receiver Operating Characteristic

## Authors’ contributions

SM, RT and GPSR collected the dataset. SM and RT preprocess the data. SM developed the classification models and implemented the algorithm. SM, AR and GPSR analyzed the results. SM and GPSR penned the manuscript. SM, AR, RT and GPSR reviewed the manuscript. All authors have read and approved the final manuscript.

## Acknowledgments

Authors are thankful to the Department of Science and Technology (DST-INSPIRE), and Council of Scientific & Industrial Research (CSIR) for fellowships and financial support, and the Department of Computational Biology, IIITD New Delhi for infrastructure and facilities. We would like to acknowledge that Figures were created using BioRender.com and chatgpt for language modification.

## References

Aoude, L. G., Xu, M., Zhao, Z. Z., Kovacs, M., Palmer, J. M., Johansson, P., Symmons, J., Trent, J. M., Martin, N. G., Montgomery, G. W., Brown, K. M., & Hayward, N. K. (2014). Assessment of PALB2 as a candidate melanoma susceptibility gene. PloS One, 9(6), e100683. 10.1371/journal.pone.0100683

Bannon, J., Cantor, C. R., & Mishra, B. (2024). A Graph Curvature-Based Pipeline for Discovering Immune Checkpoint Response Biomarkers. BioRxiv. 10.1101/2024.09.04.611306

Bhalla, S., Kaur, H., Dhall, A., & Raghava, G. P. S. (2019). Prediction and Analysis of Skin Cancer Progression using Genomics Profiles of Patients. Scientific Reports, 9(1), 15790. 10.1038/s41598-019-52134-4

Bisaccia, J., Meyer, S., Bertrand-Chapel, A., Hecquet, Q., Barbet, V., Kaniewski, B., Leon, S., Gadot, N., Rochet, I., Fajnorova, I., Leblond, P., Cordier-Bussat, M., Corradini, N., Vasiljevic, A., Billaud, M., Picard, C., Broutier, L., Gallerne, C., Dutour, A., … Castets, M. (2023). The TLR3 L412F polymorphism prevents TLR3-mediated tumor cell death induction in pediatric sarcomas. Cell Death Discovery, 9(1), 230. 10.1038/s41420-023-01513-y

Cannon, M., Stevenson, J., Stahl, K., Basu, R., Coffman, A., Kiwala, S., McMichael, J. F., Kuzma, K., Morrissey, D., Cotto, K., Mardis, E. R., Griffith, O. L., Griffith, M., & Wagner, A. H. (2024). DGIdb 5.0: rebuilding the drug-gene interaction database for precision medicine and drug discovery platforms. Nucleic Acids Research, 52(D1), D1227–D1235. 10.1093/nar/gkad1040

Chen, Y.-Y., Chang, W.-A., Lin, E.-S., Chen, Y.-J., & Kuo, P.-L. (2019). Expressions of HLA Class II Genes in Cutaneous Melanoma Were Associated with Clinical Outcome: Bioinformatics Approaches and Systematic Analysis of Public Microarray and RNA-Seq Datasets. Diagnostics (Basel, Switzerland), 9(2). 10.3390/diagnostics9020059

Christodoulou, E., Rashid, M., Pacini, C., Droop, A., Robertson, H., Groningen, T. van, Teunisse, A. F. A. S., Iorio, F., Jochemsen, A. G., Adams, D. J., & Doorn, R. van. (2021). Analysis of CRISPR-Cas9 screens identifies genetic dependencies in melanoma. Pigment Cell & Melanoma Research, 34(1), 122–131. 10.1111/pcmr.12919

Darst, B. F., Malecki, K. C., & Engelman, C. D. (2018). Using recursive feature elimination in random forest to account for correlated variables in high dimensional data. BMC Genetics, 19(Suppl 1), 65. 10.1186/s12863-018-0633-8

Ding, S., Duan, T., Yu, Y., Wu, H., Xu, Z., Yan, D., Meng, Q., Liu, Q., & Mu, Z. (2022). MYL2 is a potential diagnostic, prognostic, and immune biomarker for HNSC. Research Square. 10.21203/rs.3.rs-2142569/v1

Ding, X., Wang, W., Tao, X., Li, Z., & Huang, Y. (2023). Construction of a novel prognostic model in skin cutaneous melanoma based on chemokines-related gene signature. Scientific Reports, 13(1), 18172. 10.1038/s41598-023-44598-2

Esener, İ. I., Ergi□n, S., & YǼksel, T. (2015). A feature selection analysis in breast cancer diagnosis. 2015 Medical Technologies National Conference (TIPTEKNO), 1–4. 10.1109/TIPTEKNO.2015.7374558

Fabregat, A., Sidiropoulos, K., Viteri, G., Forner, O., Marin-Garcia, P., Arnau, V., D’Eustachio, P., Stein, L., & Hermjakob, H. (2017). Reactome pathway analysis: a high-performance in-memory approach. BMC Bioinformatics, 18(1), 142. 10.1186/s12859-017-1559-2

Fang, Y., Yu, H., & Zhou, H. (2022). MS4A15 acts as an oncogene in ovarian cancer through reprogramming energy metabolism. Biochemical and Biophysical Research Communications, 598, 47–54. 10.1016/j.bbrc.2022.01.128

Garman, B., Anastopoulos, I. N., Krepler, C., Brafford, P., Sproesser, K., Jiang, Y., Wubbenhorst, B., Amaravadi, R., Bennett, J., Beqiri, M., Elder, D., Flaherty, K. T., Frederick, D. T., Gangadhar, T. C., Guarino, M., Hoon, D., Karakousis, G., Liu, Q., Mitra, N., … Nathanson, K. L. (2017). Genetic and Genomic Characterization of 462 Melanoma Patient-Derived Xenografts, Tumor Biopsies, and Cell Lines. Cell Reports, 21(7), 1936–1952. 10.1016/j.celrep.2017.10.052

Geeitha, S., Ravishankar, K., Cho, J., & Easwaramoorthy, S. V. (2024). Integrating cat boost algorithm with triangulating feature importance to predict survival outcome in recurrent cervical cancer. Scientific Reports, 14(1), 19828. 10.1038/s41598-024-67562-0

Geng, Y., Sun, Y.-J., Song, H., Miao, Q.-J., Wang, Y.-F., Qi, J.-L., Xu, X.-L., & Sun, J.-F. (2023). Construction and Identification of an NLR-Associated Prognostic Signature Revealing the Heterogeneous Immune Response in Skin Cutaneous Melanoma. Clinical, Cosmetic and Investigational Dermatology, 16, 1623–1639. 10.2147/CCID.S410723

Geurts, P., Ernst, D., & Wehenkel, L. (2006). Extremely randomized trees. Machine Learning, 63(1), 3–42. 10.1007/s10994-006-6226-1

Guan, J., Gupta, R., & Filipp, F. V. (2015). Cancer systems biology of TCGA SKCM: efficient detection of genomic drivers in melanoma. Scientific Reports, 5, 7857. 10.1038/srep07857

Guyon, I., Weston, J., Barnhill, S., & Vapnik, V. (2002). Gene Selection for Cancer Classification using Support Vector Machines. Machine Learning, 46(1), 389–422. 10.1023/A:1012487302797

He, J., Wu, M., Xiong, L., Gong, Y., Yu, R., Peng, W., Li, L., Li, L., Tian, S., Wang, Y., Tao, Q., & Xiang, T. (2020). BTB/POZ zinc finger protein ZBTB16 inhibits breast cancer proliferation and metastasis through upregulating ZBTB28 and antagonizing BCL6/ZBTB27. Clinical Epigenetics, 12(1), 82. 10.1186/s13148-020-00867-9

He, Y., & Wang, X. (2023). Identifying biomarkers associated with immunotherapy response in melanoma by multi-omics analysis. Computers in Biology and Medicine, 167, 107591. 10.1016/j.compbiomed.2023.107591

Huang, R., Li, M., Zeng, Z., Zhang, J., Song, D., Hu, P., Yan, P., Xian, S., Zhu, X., Chang, Z., Zhang, J., Guo, J., Yin, H., Meng, T., & Huang, Z. (2022). The Identification of Prognostic and Metastatic Alternative Splicing in Skin Cutaneous Melanoma. Cancer Controli: Journal of the Moffitt Cancer Center, 29, 10732748211051554. 10.1177/10732748211051554

Ji, G., Luo, B., Chen, L., Shen, G., & Tian, T. (2021). GBP2 Is a Favorable Prognostic Marker of Skin Cutaneous Melanoma and Affects Its Progression via the Wnt/beta-catenin Pathway. Annals of Clinical and Laboratory Science, 51(6), 772–782.

Jia, G., Song, Z., Xu, Z., Tao, Y., Wu, Y., & Wan, X. (2021). Screening of gene markers related to the prognosis of metastatic skin cutaneous melanoma based on Logit regression and survival analysis. BMC Medical Genomics, 14(1), 96. 10.1186/s12920-021-00923-0

Jiang, L., Xia, Z., Zhu, R., Gong, H., Wang, J., Li, J., & Wang, L. (2023). Diabetes risk prediction model based on community follow-up data using machine learning. Preventive Medicine Reports, 35, 102358. 10.1016/j.pmedr.2023.102358

Jing Hui Nana Sun, Y. L. I. U. C. Y. U. Y. K. E. Y. C. A. O. A. Y. U. Q. K. Y. U. N. L. I. U. (2024). Bioinformatics comprehensive analysis confirmed the potential involvement of *SLC22A1* in lower-grade glioma progression and prognosis. BIOCELL, 48(5), 803–815. 10.32604/biocell.2024.047122

Kaidbey, K. H., Agin, P. P., Sayre, R. M., & Kligman, A. M. (1979). Photoprotection by melanin--a comparison of black and Caucasian skin. Journal of the American Academy of Dermatology, 1(3), 249–260. 10.1016/s0190-9622(79)70018-1

Kamiya, T., Seow, S. V., Wong, D., Robinson, M., & Campana, D. (2019). Blocking expression of inhibitory receptor NKG2A overcomes tumor resistance to NK cells. The Journal of Clinical Investigation, 129(5), 2094–2106. 10.1172/JCI123955

Kamkar, I., Gupta, S. K., Phung, D., & Venkatesh, S. (2016). Stabilizing l1-norm prediction models by supervised feature grouping. Journal of Biomedical Informatics, 59, 149–168. 10.1016/j.jbi.2015.11.012

Kardynal, A., & Olszewska, M. (2014). Modern non-invasive diagnostic techniques in the detection of early cutaneous melanoma. Journal of Dermatological Case Reports, 8(1), 1–8. 10.3315/jdcr.2014.1161

Larsson, P., Pettersson, D., Engqvist, H., Werner Ronnerman, E., Forssell-Aronsson, E., Kovacs, A., Karlsson, P., Helou, K., & Parris, T. Z. (2022). Pan-cancer analysis of genomic and transcriptomic data reveals the prognostic relevance of human proteasome genes in different cancer types. BMC Cancer, 22(1), 993. 10.1186/s12885-022-10079-4

li, rui, & Zhu, F. (2023). Deciphering the prognostic and therapeutic effects of ion channel genes in the occurrence and progression in SKCM. 10.21203/rs.3.rs-3245439/v1

Li, Y., Tian, W., Zhang, H., Zhang, Z., Zhao, Q., Chang, L., Lei, N., & Zhang, W. (2022). MSLN Correlates With Immune Infiltration and Chemoresistance as a Prognostic Biomarker in Ovarian Cancer. Frontiers in Oncology, 12, 830570. 10.3389/fonc.2022.830570

Lin, W., Tan, Z.-Y., & Fang, X.-C. (2024). Identification of m6A-related lncRNAs-based signature for predicting the prognosis of patients with skin cutaneous melanoma. SLAS Technology, 29(1), 100101. 10.1016/j.slast.2023.08.001

Liu, Y., Li, X., Yang, J., Chen, S., Zhu, C., Shi, Y., Dang, S., Zhang, W., & Li, W. (2024). Pan-cancer analysis of SLC2A family genes as prognostic biomarkers and therapeutic targets. Heliyon, 10(8), e29655. 10.1016/j.heliyon.2024.e29655

Lombard, D. B., Cierpicki, T., & Grembecka, J. (2019). Combined MAPK Pathway and HDAC Inhibition Breaks Melanoma. Cancer Discovery, 9(4), 469–471. 10.1158/2159-8290.CD-19-0069

Ma, C., & Xie, L. (2024). Prognostic model development and clinical correlation of eight key genes in skin cutaneous melanoma. Heliyon, 10(13), e33930. 10.1016/j.heliyon.2024.e33930

Ma, S., & Huang, J. (2008). Penalized feature selection and classification in bioinformatics. Briefings in Bioinformatics, 9(5), 392–403. 10.1093/bib/bbn027

Marzagalli, M., Ebelt, N. D., & Manuel, E. R. (2019). Unraveling the crosstalk between melanoma and immune cells in the tumor microenvironment. Seminars in Cancer Biology, 59, 236–250. 10.1016/j.semcancer.2019.08.002

Merighi, S., Simioni, C., Gessi, S., Varani, K., Mirandola, P., Tabrizi, M. A., Baraldi, P. G., & Borea, P. A. (2009). A(2B) and A(3) adenosine receptors modulate vascular endothelial growth factor and interleukin-8 expression in human melanoma cells treated with etoposide and doxorubicin. Neoplasia (New York, N.Y.), 11(10), 1064–1073. 10.1593/neo.09768

Niu, C., Wen, H., Wang, S., Shu, G., Wang, M., Yi, H., Guo, K., Pan, Q., & Yin, G. (2024). Potential prognosis and immunotherapy predictor TFAP2A in pan-cancer. Aging, 16(2), 1021–1048. 10.18632/aging.205225

Panda, P., & Priyadarshi, A. (2022). RFE-ACO-RF: An approach for Cancer Microarray Data Diagnosis. 2022 International Conference on Emerging Smart Computing and Informatics (ESCI), 1–5.

Pedregosa, F. and Varoquaux, G. and Gramfort, A. and Michel, V., and Thirion, B. and Grisel, O. and Blondel, M. and Prettenhofer, P., and Weiss, R. and Dubourg, V. and Vanderplas, J. and Passos, A. and, & Cournapeau, D. and Brucher, M. and Perrot, M. and Duchesnay, E. (2011). Scikit-learn: Machine Learning in {P}ython. Journal of Machine Learning Research, 12, 2825--2830.

Ping, S., Wang, S., Zhao, Y., He, J., Li, G., Li, D., Wei, Z., & Chen, J. (2022). Identification and validation of a ferroptosis-related gene signature for predicting survival in skin cutaneous melanoma. Cancer Medicine, 11(18), 3529–3541. 10.1002/cam4.4706

Scatena, C., Murtas, D., & Tomei, S. (2021). Cutaneous Melanoma Classification: The Importance of High-Throughput Genomic Technologies. Frontiers in Oncology, 11, 635488. 10.3389/fonc.2021.635488

Seitz, T., Hackl, C., Freese, K., Dietrich, P., Mahli, A., Thasler, R. M., Thasler, W. E., Lang, S. A., Bosserhoff, A. K., & Hellerbrand, C. (2021). Xanthohumol, a Prenylated Chalcone Derived from Hops, Inhibits Growth and Metastasis of Melanoma Cells. Cancers, 13(3). 10.3390/cancers13030511

Severgnini, M., D’Angio, M., Bungaro, S., Cazzaniga, G., Cifola, I., & Fazio, G. (2022). Conjoined Genes as Common Events in Childhood Acute Lymphoblastic Leukemia. Cancers, 14(14). 10.3390/cancers14143523

Siegel, R. L., Giaquinto, A. N., & Jemal, A. (2024). Cancer statistics, 2024. CA: A Cancer Journal for Clinicians, 74(1), 12–49. 10.3322/caac.21820

Sinagra, E., & Sciume, C. (2020). Ileal Melanoma, A Rare Cause of Small Bowel Obstruction: Report of a Case, and Short Literature Review. Current Radiopharmaceuticals, 13(1), 56–62. 10.2174/1874471012666191015101410

Srinivasan, S., Kalinava, N., Aldana, R., Li, Z., van Hagen, S., Rodenburg, S. Y. A., Wind-Rotolo, M., Qian, X., Sasson, A. S., Tang, H., & Kirov, S. (2021). Misannotated Multi-Nucleotide Variants in Public Cancer Genomics Datasets Lead to Inaccurate Mutation Calls with Significant Implications. Cancer Research, 81(2), 282–288. 10.1158/0008-5472.CAN-20-2151

Sun, L., Guan, Z., Wei, S., Tan, R., Li, P., & Yan, L. (2019). Identification of Long Non-coding and Messenger RNAs Differentially Expressed Between Primary and Metastatic Melanoma. Frontiers in Genetics, 10, 292. 10.3389/fgene.2019.00292

Tao, L., Cui, Y., Sun, J., Cao, Y., Dai, Z., Ge, X., Zhang, L., Ma, R., & Liu, Y. (2023). Bioinformatics-based analysis reveals elevated CYTL1 as a potential therapeutic target for BRAF-mutated melanoma. Frontiers in Cell and Developmental Biology, 11, 1171047. 10.3389/fcell.2023.1171047

Tihagam, R. D., & Bhatnagar, S. (2023). A multi-platform normalization method for meta-analysis of gene expression data. Methods (San Diego, Calif.), 217, 43–48. 10.1016/j.ymeth.2023.06.012

Tracey, E. H., & Vij, A. (2019). Updates in Melanoma. Dermatologic Clinics, 37(1), 73–82. 10.1016/j.det.2018.08.003

Wang, C., Zhu, X., Zheng, M., Chen, Y., Jiang, R., He, X., Liu, Z., Lu, Z., Wang, Z., & Yang, Y. (2024). Pan-cancer analysis of PSCA that is associated with immune infiltration and affects patient prognosis. PloS One, 19(6), e0298469. 10.1371/journal.pone.0298469

Wang, F., Zheng, A., Zhang, D., Zou, T., Xiao, M., Chen, J., Wen, B., Wen, Q., Wu, X., Li, M., Du, F., Chen, Y., Zhao, Y., Shen, J., Xiang, S., Li, J., Deng, S., Zhang, Z., Yi, T., & Xiao, Z. (2022). Molecular profiling of core immune-escape genes highlights LCK as an immune-related prognostic biomarker in melanoma. Frontiers in Immunology, 13, 1024931. 10.3389/fimmu.2022.1024931

Wang, J., & Yang, J. (2022). Identification of significant genes with a poor prognosis in skin cutaneous malignant melanoma based on a bioinformatics analysis. Annals of Translational Medicine, 10(8), 448. 10.21037/atm-22-1163

Wang, Z., & Liu, Y. (2021). MicroRNA-633 enhances melanoma cell proliferation and migration by suppressing KAI1. Oncology Letters, 21(2), 88. 10.3892/ol.2020.12349

Xiao, B., Liu, L., Li, A., Wang, P., Xiang, C., Li, H., & Xiao, T. (2021). Identification and validation of immune-related lncRNA prognostic signatures for melanoma. Immunity, Inflammation and Disease, 9(3), 1044–1054. 10.1002/iid3.468

Xiong, T.-F., Pan, F.-Q., Liang, Q., Luo, R., Li, D., Mo, H., & Zhou, X. (2020). Prognostic value of the expression of chemokines and their receptors in regional lymph nodes of melanoma patients. Journal of Cellular and Molecular Medicine, 24(6), 3407–3418. 10.1111/jcmm.15015

Yan, C., Yang, Y., Tang, Y., Zheng, X., & Xu, B. (2022). The Critical Gene Screening to Prevent Chromophobe Cell Renal Carcinoma Metastasis through TCGA and WGCNA. Journal of Oncology, 2022, 2909095. 10.1155/2022/2909095

Yan, J., Wu, X., Yu, J., Zhu, Y., & Cang, S. (2020). Prognostic Role of Tumor Mutation Burden Combined With Immune Infiltrates in Skin Cutaneous Melanoma Based on Multi-Omics Analysis. Frontiers in Oncology, 10, 570654. 10.3389/fonc.2020.570654

Yang, E., Ding, Q., Fan, X., Ye, H., Xuan, C., Zhao, S., Ji, Q., Yu, W., Liu, Y., Cao, J., Fang, M., & Ding, X. (2023). Machine learning modeling and prognostic value analysis of invasion-related genes in cutaneous melanoma. Computers in Biology and Medicine, 162, 107089. 10.1016/j.compbiomed.2023.107089

Yang, R.-H., Liang, B., Li, J.-H., Pi, X.-B., Yu, K., Xiang, S.-J., Gu, N., Chen, X.-D., & Zhou, S.-T. (2021). Identification of a novel tumour microenvironment-based prognostic biomarker in skin cutaneous melanoma. Journal of Cellular and Molecular Medicine, 25(23), 10990–11001. 10.1111/jcmm.17021

Yang, W., Ge, F., Lu, S., Shan, Z., Peng, L., Chai, J., Liu, H., Li, B., Zhang, Z., Huang, J., Hua, Y., & Zhang, Y. (2021). LncRNA MSC-AS1 Is a Diagnostic Biomarker and Predicts Poor Prognosis in Patients With Gastric Cancer by Integrated Bioinformatics Analysis. Frontiers in Medicine, 8, 795427. 10.3389/fmed.2021.795427

Zhang, C., Long, Z., Yan, C., Jalloh, M. S., Fang, Y., Bao, L., Sun, B., Shao, W., Sun, G., Fu, Y., Zhang, Q., Wang, W., Wang, L., Ren, J., Sun, Q., Tang, D., & Wang, D. (2021). Bioinformatics Analysis of ISLR in Pan-Cancer and its Correlation with Tumor Immunity. Research Square. 10.21203/rs.3.rs-970091/v1

Zhang, H., Cui, B., Zhou, Y., Wang, X., Wu, W., Wang, Z., Dai, Z., Cheng, Q., & Yang, K. (2021). B2M overexpression correlates with malignancy and immune signatures in human gliomas. Scientific Reports, 11(1), 5045. 10.1038/s41598-021-84465-6

Zia, A., Litvin, Y., Voskoboynik, R., Klein, A., & Shachaf, C. (2023). Transcriptome Analysis Identifies Oncogenic Tissue Remodeling during Progression from Common Nevi to Early Melanoma. The American Journal of Pathology, 193(7), 995–1004. 10.1016/j.ajpath.2023.03.016

