## Supplementary Table S1, Supplementary Table S2-S8, Supplementary Table S9 for "Identification of Multiple Prognostic Biomarker Sets for Risk Stratification in SKCM"

**Mailing Address of Authors**

***Corresponding Author**

Prof. Gajendra P. S. Raghava
Department of Computational Biology
Indraprastha Institute of Information Technology, Delhi
Okhla Industrial Estate, New Delhi, India – 110020


Website: <http://webs.iiitd.edu.in/raghava/>

**Supplementary Table S1**: **The table shows the summary of Previous Studies on Prognostic Biomarkers in SKCM**

| **S. No** | **Study Name** | **Database** | **Analysis** | **Specific Study type** | **Statistical Method** | **ML Model** | **PMID** |
| --- | --- | --- | --- | --- | --- | --- | --- |
| 1 | Machine learning modeling and prognostic value analysis of  invasion-related genes in cutaneous melanoma | GEO,TCGA & CancerSEA | DEGS, Survival Aanlysis, Enrichment Analysis,Lasso | Invasion associated genes | Yes | Yes | 37267825 |
| 2 | Identification of m6A-related lncRNAs-based signature for predicting the prognosis of patients with skin cutaneous melanoma | TCGA, GEO, lncbase,miRBase and TargetScan | Univariate & Pearson Correlation,Enrichment Analysis,Lasso | m6A related LncRNAs | Yes | No | 37541541 |
| 3 | Skin cutaneous melanoma properties of immune‐related lncRNAs  identifying potential prognostic biomarkers | TCGA,GTEx & immLnc | WGCNA, DEGS, Survival Aanlysis, Enrichment Analysis,Lasso | immune related LncRNAs | Yes | No | 35361740 |
| 4 | Identification and validation of a ferroptosis-related gene signature for predicting survival in skin cutaneous  melanoma | TCGA, GEO, and GTEx. | Univariate, multivariate,Enrichment Analysis,Lasso | Ferroptosis related gene signature | Yes | No | 35373463 |
| 5 | Construction and Identification of an  NLR-Associated Prognostic Signature Revealing  the Heterogeneous Immune Response in Skin  Cutaneous Melanoma | TCGA & GEO | Univariate Analysis,Lasso | NLR-associated | Yes | No | 37396711 |
| 6 | Identification and validation of  ferroptosis-related lncRNA  signature as a prognostic model  for skin cutaneous melanoma | TCGA ,GEO & Ensembl | DEGS, Survival Aanlysis,Lasso | Ferroptosis related lncRNAs signature | Yes | No | 36248853 |
| 7 | New Prognostic Biomarkers and  Drug Targets for Skin Cutaneous  Melanoma via Comprehensive  Bioinformatic Analysis and Validation | TCGA & GEO | DEGS, WGCNA,GWAS,Survival & Enrichment Analysis,Lasso | 10 hub genes | Yes | No | 34722301 |
| 8 | Identification of Novel Molecular  Therapeutic Targets and Their  Potential Prognostic Biomarkers  Based on Cytolytic Activity in  Skin Cutaneous Melanoma | TCGA & GEO | TME, Survival, Enrichment Analysis,Lasso | Cyt related genes | Yes | No | 35345444 |
| 9 | Identification and validation of immune‐related lncRNA  prognostic signatures for melanoma | TCGA | Survival Analysis,Lasso | immune related LncRNAs | Yes | No | 34077998 |
| 10 | Machine learning-derived identifcation  of tumor-infltrating immune cell-related  signature for improving prognosis  and immunotherapy responses in patients  with skin cutaneous melanoma | TCGA & GEO | WGCNA, ICI, Survival Aanlysis, Enrichment Analysis,Lasso | Tumor infiltrating immune cell related | Yes | Yes | 37752452 |
| 11 | Identification of Pyroptosis-Relevant Signature in Tumor  Immune Microenvironment and Prognosis in Skin Cutaneous  Melanoma Using Network Analysis | TCGA & GEO | WGCNA, Statistical Analysis,Lasso | Pyroptopsis | Yes | No | 36818162 |
| 12 | Development and validation of an immune  gene set-based prognostic signature in  cutaneous melanoma | TCGA & GEO | DEGS, Enrichement Analysis, Survival Analysis,Lasso | Immune related geneset | Yes | No | 34291650 |
| 13 | Prognostic model development and clinical correlation of eight key genes in skin cutaneous melanoma | TCGA | Univaraite Analysis,Lasso | Eight key genes | Yes | No | 39071565 |
| 14 | The predictive efficacy of programmed cell death in immunotherapy of melanoma: A comprehensive analysis of gene expression data for programmed cell death biomarker and therapeutic target discovery | Xena- browser(TCGA) | Survival Analysis,Lasso | PCD related | Yes | No | 38140739 |
| 15 | Characteristics and significance of programmed cell death-related gene expression signature in skin cutaneous melanoma | TCGA | Univaraite , Enrichment & PPI analysis,Lasso | PCD | Yes | No | 38766879 |

**Supplementary Table S2: The table shows the performance on 20 features selected using selectkbest approach.**

|  | **Training** | | | | | | **Testing** | | | | | |
| --- | --- | --- | --- | --- | --- | --- | --- | --- | --- | --- | --- | --- |
| **Model** | **Acc** | **AUC** | **Sens** | **Spec** | **MCC** | **Kappa** | **Acc** | **AUC** | **Sens** | **Spec** | **MCC** | **Kappa** |
| **RF** | **0.61** | **0.82** | **0.61** | **0.60** | **0.48** | **0.48** | **0.51** | **0.82** | **0.51** | **0.50** | **0.35** | **0.35** |
| SVM | 0.59 | 0.78 | 0.59 | 0.57 | 0.45 | 0.45 | 0.58 | 0.80 | 0.58 | 0.56 | 0.44 | 0.44 |
| LR | 0.45 | 0.74 | 0.45 | 0.42 | 0.27 | 0.27 | 0.42 | 0.71 | 0.42 | 0.42 | 0.23 | 0.23 |
| **ET** | **0.61** | **0.84** | **0.61** | **0.60** | **0.48** | **0.48** | **0.53** | **0.82** | **0.53** | **0.53** | **0.38** | **0.38** |
| XGB | 0.60 | 0.80 | 0.60 | 0.59 | 0.46 | 0.46 | 0.58 | 0.81 | 0.58 | 0.59 | 0.44 | 0.44 |

**Supplementary Table S3**. **The table shows the performance on 50 features selected using selectkbest approach.**

|  | **Training** | | | | | | **Testing** | | | | | |
| --- | --- | --- | --- | --- | --- | --- | --- | --- | --- | --- | --- | --- |
| **Model** | **Acc** | **AUC** | **Sens** | **Spec** | **MCC** | **Kappa** | **Acc** | **AUC** | **Sens** | **Spec** | **MCC** | **Kappa** |
| **RF** | **0.66** | **0.86** | **0.66** | **0.67** | **0.56** | **0.55** | **0.66** | **0.86** | **0.66** | **0.67** | **0.55** | **0.55** |
| SVM | 0.65 | 0.86 | 0.65 | 0.66 | 0.54 | 0.53 | 0.66 | 0.86 | 0.66 | 0.65 | 0.55 | 0.55 |
| LR | 0.57 | 0.77 | 0.57 | 0.56 | 0.43 | 0.42 | 0.51 | 0.75 | 0.51 | 0.49 | 0.36 | 0.35 |
| **ET** | **0.68** | **0.88** | **0.68** | **0.68** | **0.58** | **0.58** | **0.69** | **0.88** | **0.69** | **0.69** | **0.59** | **0.59** |
| XGB | 0.67 | 0.84 | 0.67 | 0.67 | 0.56 | 0.56 | 0.61 | 0.85 | 0.61 | 0.60 | 0.49 | 0.48 |

**Supplementary Table S4: The table shows the performance on 100 features selected using selectkbest approach.**

|  | **Training** | | | | | | **Testing** | | | | | |
| --- | --- | --- | --- | --- | --- | --- | --- | --- | --- | --- | --- | --- |
| **Model** | **Acc** | **AUC** | **Sens** | **Spec** | **MCC** | **Kappa** | **Acc** | **AUC** | **Sens** | **Spec** | **MCC** | **Kappa** |
| **RF** | **0.68** | **0.87** | **0.68** | **0.70** | **0.58** | **0.58** | **0.69** | **0.86** | **0.69** | **0.69** | **0.59** | **0.59** |
| **SVM** | **0.66** | **0.87** | **0.66** | **0.68** | **0.55** | **0.55** | **0.69** | **0.89** | **0.69** | **0.70** | **0.59** | **0.59** |
| LR | 0.66 | 0.81 | 0.66 | 0.67 | 0.56 | 0.55 | 0.63 | 0.79 | 0.63 | 0.60 | 0.50 | 0.50 |
| **ET** | **0.71** | **0.88** | **0.71** | **0.72** | **0.62** | **0.61** | **0.66** | **0.86** | **0.66** | **0.64** | **0.55** | **0.55** |
| XGB | 0.66 | 0.86 | 0.66 | 0.67 | 0.55 | 0.55 | 0.68 | 0.86 | 0.68 | 0.68 | 0.58 | 0.58 |

**Supplementary Table S5**: **The table shows the performance on 50 features selected by Support vector classifier- L1 regularization (SVC-L1) based on Feature selection approach.**

|  | **Training** | | | | | | **Testing** | | | | | |
| --- | --- | --- | --- | --- | --- | --- | --- | --- | --- | --- | --- | --- |
| **Model** | **Acc** | **AUC** | **Sens** | **Spec** | **MCC** | **Kappa** | **Acc** | **AUC** | **Sens** | **Spec** | **MCC** | **Kappa** |
| **RF** | **0.72** | **0.91** | **0.72** | **0.73** | **0.63** | **0.62** | **0.70** | **0.88** | **0.70** | **0.71** | **0.61** | **0.61** |
| **ET** | **0.82** | **0.95** | **0.82** | **0.84** | **0.76** | **0.75** | **0.68** | **0.89** | **0.68** | **0.70** | **0.58** | **0.58** |
| XGB | 0.68 | 0.89 | 0.68 | 0.68 | 0.58 | 0.58 | 0.64 | 0.82 | 0.64 | 0.64 | 0.52 | 0.52 |
| KNN | 0.69 | 0.88 | 0.69 | 0.73 | 0.60 | 0.59 | 0.59 | 0.80 | 0.59 | 0.59 | 0.48 | 0.45 |
| LightGBM | 0.75 | 0.92 | 0.75 | 0.77 | 0.68 | 0.67 | 0.72 | 0.86 | 0.72 | 0.71 | 0.62 | 0.62 |
| GB | 0.64 | 0.85 | 0.64 | 0.65 | 0.52 | 0.52 | 0.60 | 0.83 | 0.60 | 0.60 | 0.48 | 0.47 |
| Adaboost | 0.70 | 0.87 | 0.70 | 0.73 | 0.61 | 0.60 | 0.65 | 0.82 | 0.65 | 0.67 | 0.54 | 0.53 |
| **Catboost** | **0.68** | **0.92** | **0.68** | **0.68** | **0.58** | **0.57** | **0.66** | **0.91** | **0.66** | **0.67** | **0.56** | **0.55** |

**Supplementary Table S6:** **The table shows the performance on 20 features selected using Recursive Feature Elimination approach.**

|  | **Training** | | | | | | **Testing** | | | | | |
| --- | --- | --- | --- | --- | --- | --- | --- | --- | --- | --- | --- | --- |
| **Model** | **Acc** | **AUC** | **Sens** | **Spec** | **MCC** | **Kappa** | **Acc** | **AUC** | **Sens** | **Spec** | **MCC** | **Kappa** |
| RF | 0.69 | 0.89 | 0.69 | 0.69 | 0.60 | 0.59 | 0.69 | 0.85 | 0.69 | 0.69 | 0.59 | 0.59 |
| ET | 0.74 | 0.92 | 0.74 | 0.75 | 0.66 | 0.66 | 0.70 | 0.87 | 0.70 | 0.71 | 0.61 | 0.61 |
| XGB | 0.68 | 0.87 | 0.68 | 0.69 | 0.58 | 0.58 | 0.64 | 0.83 | 0.64 | 0.65 | 0.52 | 0.52 |
| KNN | 0.63 | 0.85 | 0.63 | 0.65 | 0.52 | 0.51 | 0.55 | 0.77 | 0.55 | 0.53 | 0.40 | 0.39 |
| LightGBM | 0.70 | 0.90 | 0.70 | 0.71 | 0.61 | 0.61 | 0.60 | 0.85 | 0.60 | 0.61 | 0.47 | 0.54 |
| GB | 0.60 | 0.81 | 0.60 | 0.61 | 0.47 | 0.47 | 0.50 | 0.76 | 0.50 | 0.51 | 0.34 | 0.42 |
| Adaboost | 0.56 | 0.82 | 0.56 | 0.64 | 0.42 | 0.41 | 0.45 | 0.74 | 0.45 | 0.54 | 0.28 | 0.27 |
| **Catboost** | **0.66** | **0.89** | **0.66** | **0.67** | **0.56** | **0.55** | **0.72** | **0.90** | **0.72** | **0.71** | **0.63** | **0.62** |

**Supplementary Table S7: The table shows the performance on 50 features selected using Recursive Feature Elimination approach.**

|  | **Training** | | | | | | **Testing** | | | | | |
| --- | --- | --- | --- | --- | --- | --- | --- | --- | --- | --- | --- | --- |
| **Model** | **Acc** | **AUC** | **Sens** | **Spec** | **MCC** | **Kappa** | **Acc** | **AUC** | **Sens** | **Spec** | **MCC** | **Kappa** |
| RF | 0.71 | 0.91 | 0.71 | 0.72 | 0.62 | 0.62 | 0.65 | 0.86 | 0.65 | 0.63 | 0.53 | 0.53 |
| ET | 0.80 | 0.94 | 0.80 | 0.80 | 0.74 | 0.74 | 0.66 | 0.89 | 0.66 | 0.66 | 0.55 | 0.55 |
| XGB | 0.71 | 0.90 | 0.71 | 0.71 | 0.62 | 0.61 | 0.67 | 0.86 | 0.67 | 0.67 | 0.56 | 0.56 |
| KNN | 0.67 | 0.89 | 0.67 | 0.70 | 0.58 | 0.56 | 0.52 | 0.79 | 0.52 | 0.50 | 0.38 | 0.36 |
| LightGBM | 0.74 | 0.92 | 0.74 | 0.76 | 0.66 | 0.66 | 0.63 | 0.86 | 0.63 | 0.63 | 0.51 | 0.50 |
| GB | 0.60 | 0.82 | 0.60 | 0.60 | 0.47 | 0.46 | 0.59 | 0.80 | 0.59 | 0.58 | 0.46 | 0.45 |
| **Catboost** | **0.68** | **0.91** | **0.68** | **0.68** | **0.59** | **0.58** | **0.65** | **0.90** | **0.65** | **0.64** | **0.54** | **0.53** |

**Supplementary Table S8: The table shows the performance on 20 features using Sequential feature selection approach.**

|  | **Training** | | | | | | **Testing** | | | | | |
| --- | --- | --- | --- | --- | --- | --- | --- | --- | --- | --- | --- | --- |
| **Model** | **Acc** | **AUC** | **Sens** | **Spec** | **MCC** | **Kappa** | **Acc** | **AUC** | **Sens** | **Spec** | **MCC** | **Kappa** |
| RF | 0.67 | 0.88 | 0.67 | 0.68 | 0.57 | 0.56 | 0.63 | 0.84 | 0.63 | 0.63 | 0.50 | 0.50 |
| ET | 0.73 | 0.90 | 0.73 | 0.73 | 0.64 | 0.64 | 0.60 | 0.85 | 0.60 | 0.61 | 0.47 | 0.47 |
| XGB | 0.65 | 0.85 | 0.65 | 0.65 | 0.53 | 0.53 | 0.55 | 0.82 | 0.55 | 0.55 | 0.40 | 0.39 |
| KNN | 0.61 | 0.83 | 0.61 | 0.61 | 0.48 | 0.47 | 0.57 | 0.78 | 0.57 | 0.54 | 0.43 | 0.42 |
| LightGBM | 0.68 | 0.88 | 0.68 | 0.68 | 0.57 | 0.57 | 0.57 | 0.82 | 0.57 | 0.56 | 0.42 | 0.42 |
| GB | 0.58 | 0.80 | 0.58 | 0.58 | 0.44 | 0.44 | 0.52 | 0.76 | 0.52 | 0.51 | 0.37 | 0.36 |
| **Catboost** | **0.70** | **0.90** | **0.70** | **0.69** | **0.60** | **0.59** | **0.63** | **0.85** | **0.63** | **0.61** | **0.50** | **0.50** |

**Supplementary Table S9: The table shows the lists of drugs that are found to targets genes**

| **Gene** | **Drug** | **Regulatory Approval** | **Indication** | **Interaction score** |
| --- | --- | --- | --- | --- |
| **KLRC1** | MONALIZUMAB | Not Approved | antineoplastic agent | 105.01 |
| **SLC2A5** | GLUFOSFAMIDE | Not Approved |  | 3.28 |
| **SLC2A5** | STREPTOZOCIN | Approved |  | 1.141 |
| **MCOLN2** | PRU-10 | Not Approved |  | 2.91 |
| **MCOLN2** | ML SA1 | Not Approved |  | 2.91 |
| **MCOLN2** | ESTRADIOL 3-METHYL ETHER | Not Approved |  | 2.917 |
| **MCOLN2** | PRU-12 | Not Approved |  | 2.917 |
| **MCOLN2** | PHOSPHATIDYL (3,5) INOSITOL BISPHOSPHATE | Not Approved |  | 1.75 |
| **KLRC1** | MONALIZUMAB | Not Approved | antineoplastic agent | 105.01 |
| **SLC2A5** | GLUFOSFAMIDE | Not Approved |  | 3.28 |
| **SLC2A5** | STREPTOZOCIN | Approved |  | 1.141 |
| **MCOLN2** | ML2-SA1 | Not Approved |  | 8.75 |
| **B2M** | GVAX | Not Approved |  | 13.12 |
| **B2M** | BORTEZOMIB | Approved | antineoplastic agent | 0.15 |
| **B2M** | IPILIMUMAB | Approved | antineoplastic agent | 0.65 |
| **B2M** | NIVOLUMAB | Approved | antineoplastic agent | 0.43 |
| **B2M** | PEMBROLIZUMAB | Approved |  | 0.75 |
